# A regulatory TRIF/IL-1R1 axis controls T-dependent IgA production in the intestines

**DOI:** 10.64898/2026.05.22.725254

**Authors:** Chi-Chang Sung, Nimat Gafar Badmus, Michelle Shao, Christhalia Jusuf, Claire Tang, Nathalie Groot, Shin-Heng Chiou, Steven S An, Arnold B. Rabson, Qi Yang, Gaetan Barbet

## Abstract

Mucosal IgA is critical for controlling the microbiota and preventing pathogenic infection of the epithelium. The innate signals that regulate the generation of IgA remain poorly defined. Here, we identified TRIF and IL-1R1 as immune checkpoints for intestinal IgA production. In the absence of infection, *Trif^−/−^* and *Il1r1^−/−^* mice exhibited markedly elevated stool IgA and IgA-bound commensals. We show that IL-1R1 restricts IgA class-switching in a B cell-intrinsic manner, while TRIF acts extrinsically of B cells and shapes the Peyer’s patch microenvironment. Loss of either *Trif* or *Il1r1* enhanced retinoic acid metabolism and allowed premature Ccnd3 upregulation in naive B cells, favoring both IgA class-switching and differentiation into germinal center B cells. During oral vaccination, the absence or blockade of the TRIF/IL-1R1 pathway increased antigen-specific IgA production without affecting seric antigen-specific IgG levels. These findings unveil novel and local signaling targets to promote robust antigen-specific mucosal immunity.

## INTRODUCTION

The most common port of entry of pathogens are mucosal surfaces^1^, with respiratory and enteric diseases remaining a public health challenge^2^: diarrheal diseases alone cause an estimated 1.17 million deaths worldwide^3^. Distinct antibody isotypes confer specialized protection against infections in different anatomical contexts. IgG is the predominant antibody in circulating in the blood, whereas IgA is the dominant isotype at mucosal surfaces. At mucosal sites and especially in the intestines, the immune system not only preserves barrier integrity but also facilitates host–microbiota mutualism. Disruption of this finely-tuned balance can contribute to chronic inflammatory diseases such as inflammatory bowel disease (IBD)^4,5^. Through its direct influence on the microbiome, IgA serves as a pivotal gatekeeper of human health and homeostasis. Moreover, effective defense against many intestinal pathogens and toxins relies heavily on secretory IgA-mediated immune exclusion at the gut mucosa, underscoring the critical importance of mucosal IgA for protection against enteric infection^6–8^.

Despite the crucial role of IgA antibodies in barrier defense only a small fraction of licensed vaccines are delivered via mucosal routes, and there are a limited number of FDA-approved mucosal vaccines when compared with injectable vaccines^9,10^. Parenteral vaccines induce robust protection through systemic IgG production but often fail to elicit strong IgA responses at the port of entry for most pathogens^11–13^. Hence, parenteral vaccines protect from severe forms of infectious diseases, but their ability to prevent mucosal infection is often limited. There are numerous challenges to the establishment of robust and long term immune protection in the intestines: 1) mucosal immunity is biased toward tolerance to avoid overt inflammation and immune reactions against the vast load of innocuous food antigens and commensal microbes encountered daily^14^; 2) antigens administered to the gastrointestinal tract often face rapid loss of bioavailability due to gastric acidity, digestive enzymes, bile salts, mucus and epithelial barriers, reducing vaccine efficacy by weakening antigen immunogenicity^15,16^; and 3) adjuvants used in parenteral vaccines are largely ineffective at mucosal sites^17^.

In absence of a consistently effective vaccine strategy for mucosal sites, live attenuated organisms remain the most effective platform for inducing mucosal protection, exemplified by anti-*Vibrio Cholerae* or the anti-Rotavirus vaccines (Vaxchora and RotaTeq)^9^, withadditional candidates in clinical development^18^. However, live vaccines face limited suitability because of ongoing concerns about potential pathogenic reversion,^19,20^ as well as a growing population of immunosuppressed individuals who cannot be vaccinated with live vaccines^21^. These factors have motivated efforts to identify “viability signals” that recapitulate key immunostimulatory features of live microbes. Peyer’s patches (PPs) are specialized secondary lymphoid tissues embedded in the small intestine. PPs are constantly exposed to dietary and microbiota-derived antigens and thus have constant ongoing germinal center (GC) activity^22^. Notably, PPs serve as major inductive sites for intestinal IgA responses and the generation of IgA-committed B cells^22,23^. At mucosal sites, secretory IgAs are produced continuously in the absence of inflammation or infection, providing frontline protection by neutralizing pathogens and toxins at the barrier and by shaping interactions with commensal bacteria. While various factors—such as TGF-β, BAFF, IL-6, IL-21, and retinoic acid (RA)—are known to promote IgA production^24^, translating this knowledge into effective mucosal adjuvants or vaccines remains an ongoing challenge. For example, RA is known to promote intestinal IgA^25,26^ production, but also causes adverse effects^27–30^ both in the skin and the intestines.

These limitations in mucosal vaccines have prompted a growing emphasis on strategies that deliberately enhance mucosal immunity^31^. In the context of SARS-CoV-2, higher mucosal IgA has been linked to reduced risk of breakthrough infection and to improved host control, supporting its relevance as a protective correlate^32–35^. One major approach to bridge this gap is the “prime-and-pull” strategy, which pairs systemic antigenic priming with chemokine-mediated recruitment of effector cells into mucosal tissues^36–38^. This innovative approach which showed promising results in several mouse models of infection (e.g., HSV, E. coli, influenza), highlights an existing gap of knowledge in mucosal immunity.

Recently multiple studies have shown how IL-1β^39,40^ or TRIF^41^ (the mouse gene name is *Ticam1*) signaling are crucial for T follicular helper cell (T_FH_) development, GC formation, and protective antibody production. More specifically, we previously identified that the TRIF/IL-1β axis is triggered upon live, but not heat-killed, bacterial immunization, and this triggering enhanced T_FH_ differentiation, GC formation, and long-term protection through the production of IgG^42^. Since mucosal tissues are continuously exposed to microbial components from live commensals, we wanted to investigate the role of the TRIF/IL-1β pathway on IgA secretion. Counterintuitively, our observations established the role of the adaptor protein TRIF and the interleukine-1 receptor 1 (IL-1R1) as innate checkpoints of mucosal IgA production. Through cell-specific knockouts, adoptive transfers, single-cell transcriptomic analysis and a vaccination model, we will demonstrate that the TRIF/IL-1β axis acts as a brake for high-affinity IgA production. Furthermore, we show that in C57BL/6J mice (wild-type, WT) IL-1 blockade synergizes with RA signaling to improve antigen-specific IgA production. These findings highlight that innate inflammatory pathways are targetable context-dependent mechanisms that can be manipulated to enhance humoral immunity.

## RESULTS

### Both *Trif* and *Il1r1* deficiencies promote steady-state intestinal IgA but not IgG

To assess the impact of TRIF or IL-1R1 on intestinal antibody production in the absence of infection, we quantified the IgA levels in the stool and blood, as well as the IgG levels in the blood for comparison in our WT, *Trif^LPS^*^2^ (referred to as *Trif⁻^/^⁻* below) and *Il1r1⁻^/^⁻* mice weekly. We stained fecal samples from specific-pathogen-free (SPF) mice for IgA and confirmed that only a fraction of intestinal bacteria are coated by IgA, as determined by flow cytometry, as previously established^43^. Stool samples from both *Trif*- and *Il1r1*-deficient mice exhibited a higher percentage of IgA-positive bacteria over time (Fig 1A and 1B) and remained consistently higher in both male and female mice (Fig S1A). Free IgA levels in the stool, as determined by ELISA, also showed a significant increase in both *Trif*- and *Il1r1*-deficient mice compared to WT mice (Fig 1C, Fig S1B). However, IgA levels in the serum were relatively comparable between the three mouse groups (Fig 1D, Fig S1C). Furthermore, the levels of total IgG in the stools and sera were similar between the groups (Fig 1E, F, Fig S1D,E). Taken together, these observations suggest that the TRIF/IL-1R1 axis exerts a negative regulatory effect specifically on intestinal IgA and not IgG production at steady state.

**Figure 1.**
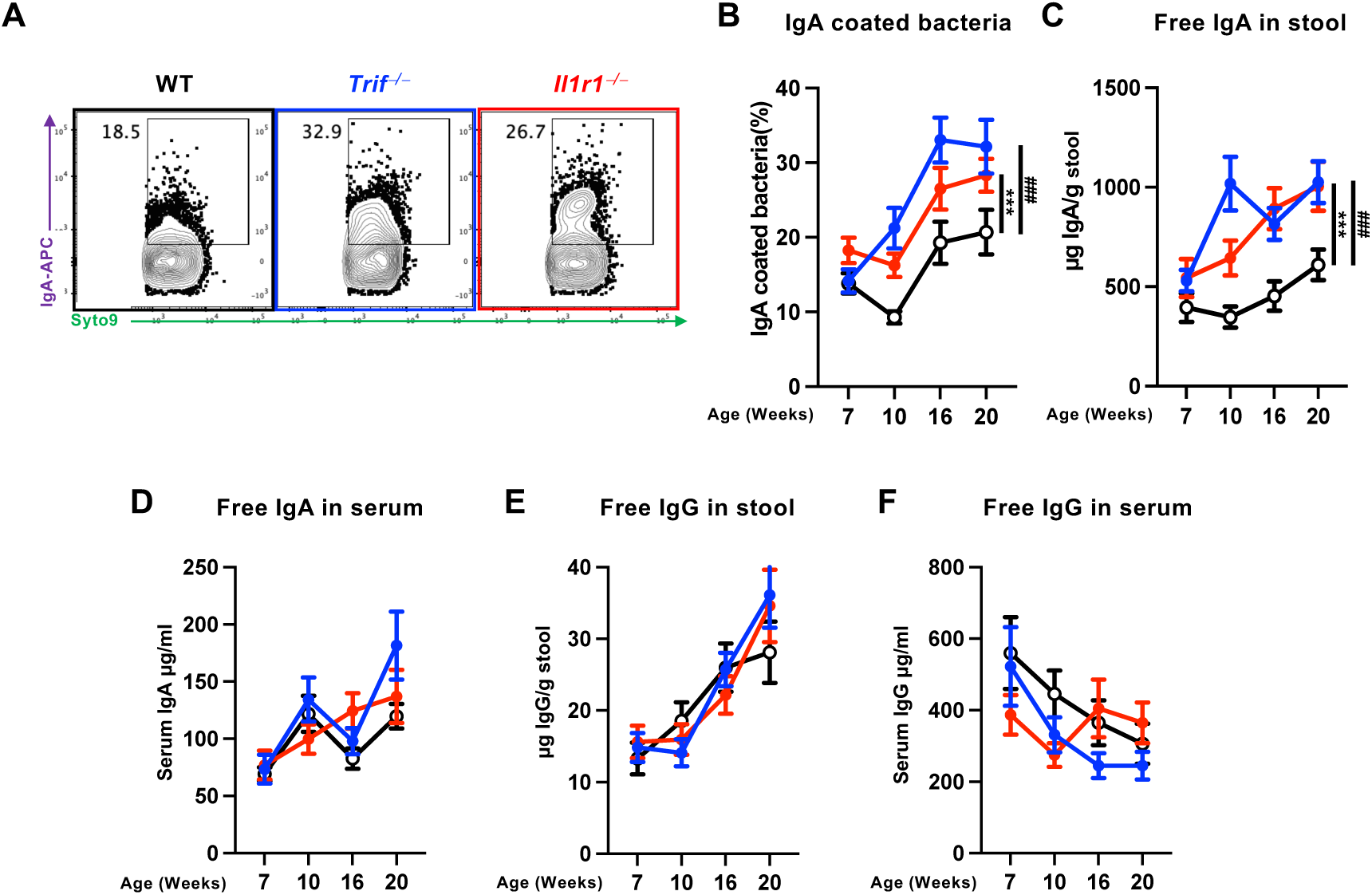
Elevated intestinal IgA but not IgG levels in *Trif^−/−^* and *Il1r1^−/−^* mice under steady-state conditions. (**A, B**) IgA-coated bacteria in stool samples from mice maintained under specific pathogen-free (SPF) conditions. (**A**) Representative flow cytometry histograms. (**B**) Percentage of IgA-coated bacteria over time. (**C, D**) Free IgA levels in stool (**C**) and serum (**D**), measured by ELISA. (**E, F**) Free IgG levels in stool (**E**) and serum (**F**), measured by ELISA. Statistical analysis: Student’s t-test; * p<0.05; ** p<0.01; *** p<0.001. *#*: Wt to *Trif ^⎻/⎻^*, *: Wt to *Il1r1^⎻/⎻^*. (n=25-29 mice per group pooled from four experiments).

We then sought to determine whether these phenotypes from *Trif* or *Il1r1* deficiencies could result in altered intestinal inflammatory status or barrier dysfunction. First, we quantified fecal Lipocalin-2 (LCN-2), a sensitive marker of intestinal inflammation. Consistent with previous reports^44^, mice infected with *Citrobacter rodentium* exhibited robust LCN-2 elevation and severe inflammation. In contrast, LCN-2 remained nearly undetectable in the stools of WT, *Trif⁻^/^⁻,* and *Il1r1⁻^/^⁻* mice in absence of infection, indicating an absence of intestinal inflammation in steady state (Fig S1F). Concomitantly, histological analysis revealed baseline levels of leukocyte infiltration, normal muscularis thickness, and preserved epithelial integrity across all groups (Fig S1G). The number of PPs per mouse was similar regardless of *Trif/Il1r1* genetic state (Fig S1H). Furthermore, intestinal permeability assays using oral FITC-dextran administration showed comparable serum FITC levels across the cohorts at all time points, indicating that the intestinal barrier remains intact in the absence of TRIF or IL-1R1 signaling (Fig S1I). Additionally, expression levels of antimicrobial peptides across different intestinal segments suggested a comparable mucosal environment (Fig S1K). Furthermore, in the PPs, the mRNA levels of T-B crosstalk-associated genes, including *Il6* and *Tgfb1*, were similar among the indicated strains (Fig S1J), suggesting that additional factors may contribute to the enhanced IgA production observed in these mice. Collectively, these findings show that *Trif* and *Il1r1* deficiencies do not induce intestinal inflammation. Therefore, the increased IgA production in these knock-out (KO) mice is not a sequela of intestinal inflammation or microanatomical defects.

### *Trif* and *Il1r1* deficiencies increased in T-dependent IgA production from the Peyer’s patches

To uncover the cellular mechanisms underlying the enhanced intestinal IgA production in *Trif⁻/⁻* and *Il1r1⁻/⁻* mice, we systematically examined the key immune populations in PPs, including dendritic cells (DC), T follicular helper (T_FH_) cells, IgA-switched B cells and plasma cells. Given that PP-resident DCs are critical for inducing gut-homing receptors and IgA switching^25,26,45–47^, we first investigated whether DC-intrinsic signaling was required. We generated DC-specific *Il1r1*-deficient mice (*Il1r1ᶠ_ˡ_/ᶠ_ˡ_ Itgax-Cre*; referred to as *Il1r1*^ΔCD11c^). However, *Il1r1*^ΔCD11c^ mice exhibited IgA-coated bacteria ratios and free fecal IgA levels comparable to those in the control *Il1r1ᶠ_ˡ_/ᶠ_ˡ_* cohort (Fig. S2A–C), demonstrating that DC-intrinsic IL-1R1 signaling is dispensable for this phenotype.

To elucidate the activation status of T cells in the KO mice, we analyzed the percentages of different T cell subsets within the two main gut-associated lymphoid tissues, the mesenteric lymph nodes (MLNs) and the PPs. Within the CD4 T cell compartment, we did not observe significant differences in the frequencies of Treg or T_H_17 cells in the intestines (Fig. S3A–D). However, we detected a significantly higher proportion of T_FH_ cells—defined as PD-1⁺CXCR5⁺ cells—within PPs from both knockout strains compared with WT mice (Fig. 2A, 2B). The selective expansion of T_FH_ cells within PPs suggested that enhanced IgA production in *Trif-* and *Il1r1*-deficient mice is associated with augmented T cell–dependent responses within GCs rather than broad alterations in intestinal T cell homeostasis.

**Figure 2.**
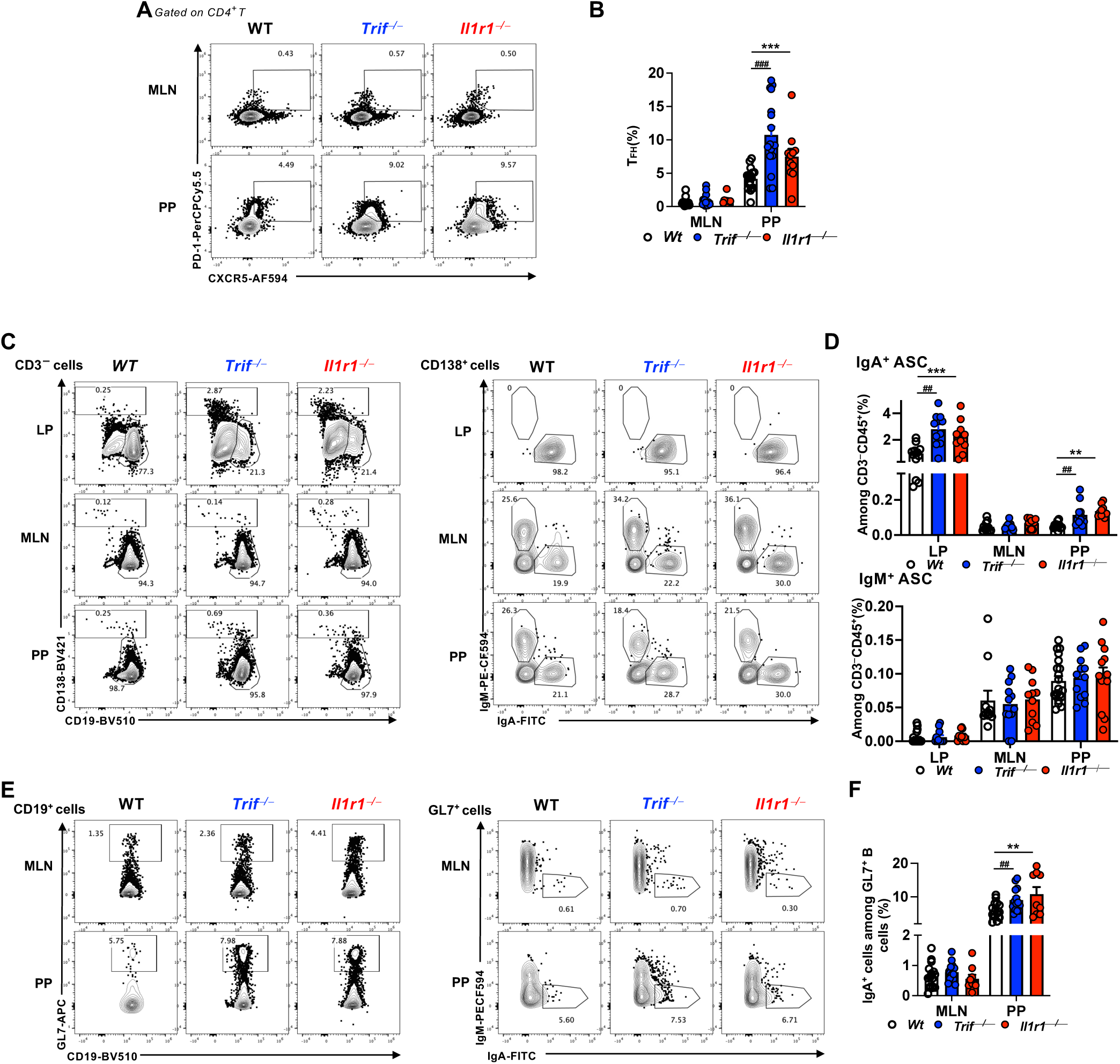
*Trif* or *Il1r1* deficiency promotes intestinal IgA^+^ B cell and T_FH_ differentiation at steady state. (A, B) Dot plots (A) and percentages (B) of CXCR5⁺PD-1⁺ CD4 T follicular helper (T_FH_) cells. (C) Flow cytometry dot plots of CD138⁺ antibody-secreting cells (ASCs) among CD3⁻CD45⁺ cells and the ratio of IgM/IgA+ cells among those ASCs. (D) and percentages of IgM/IgA+ ASCs among CD3⁻CD45⁺ cells. (E, F) Dot plots (E) and percentages (F) of IgA⁺GL7⁺ B cells among CD19⁺ cells. LP: lamina propria of the small intestine; MLN: mesenteric lymph nodes; PP: Peyer’s patches. Statistical analysis: Student’s t-test; * p<0.05; ** p<0.01; *** p<0.001. *#*: Wt to *Trif ^⎻/⎻^*, *: Wt to *Il1r1^⎻/⎻^*. Each dot represents one mouse (n = 10-12 pooled from three independent experiments).

We then assessed the levels of IgA-switched B cells and IgA producing plasma cells in both *Trif*- and *Il1r1*-deficient mice. We isolated leukocytes from the lamina propria of the small intestines (LP), MLNs, and PPs, and analyzed the composition of B cells using flow cytometry. Using the markers CD138 and CD19 to verify the ratio of antibody secreting cells (ASC, defined by CD138^+^) and conventional B cells (CD138^−^CD19^+^), we found both *Trif*- and *Il1r1*-deficient mice exhibited a higher ratio of ASC in the LP and PP when compared to WT mice (Fig 2C). Further analysis revealed an increased proportion of IgA-positive ASCs among CD45^+^ cells in KO mice. In contrast, the IgM-positive ASC remained similar between the three groups of mice (Fig 2D). Among the population of CD19^+^ B cells, we observed a higher ratio of GL7^+^ cells in PPs of both KO mice, suggesting that the *Trif*- and *Il1r1*-deficiencies facilitate GC B cell differentiation specifically within PPs (Fig 2E). Concomitantly, we noticed a higher ratio of IgA^+^ cells among GC B cells (Fig 2F) in PPs of both KO mice compared to WT mice.

Collectively, the observations we made in ASCs, GC B cells, and T_FH_ cells support a model that TRIF/IL-1R1 signaling restrains PP-driven, T cell–dependent (TD) IgA responses. However, intestinal IgAs can also be generated via T cell–independent (TI) pathways, so we next assessed the contribution of TI mechanisms using genetic disruption of TI IgA induction. We crossed *Il1r1^−/−^* mice with *Tnfrsf13b^−/−^* (TACI-deficient) mice, which exhibit impaired TI IgA responses^48^, and assessed if the level intestinal IgA level in *Il1r1* deficiency. Notably, the elevated, steady-state intestinal IgA levels driven by *Il1r1* deficiency persisted in the TACI-deficient mice (Fig. S3E-G), indicating that the IL-1R1 axis restrains IgA responses predominantly through the TD pathway.

To rule out that differences in genetic background and/or commensal microbiota among the independently maintained KO and WT strains could have contributed to the observed phenotypes, we backcrossed both KO strains to WT mice, generated F1 heterozygous mice, and intercrossed the F1 mice to obtain littermate controls for both strains of mice. This strategy allowed mice of different genotypes to share a similar genetic background and to be cohoused before weaning, thereby minimizing microbiota-driven differences. Compared to their respective littermate controls, both *Trif*- and *Il1r1*-deficient mice displayed higher IgA-bound bacteria (Fig. S4A), higher T_FH_ differentiation in their PPs but not their MLNs, (Fig. S4B and 4C) and higher percentages of ASC in the lamina propria of their small intestines and their PPs (Fig. S4D and 4E). These results, taken together, suggest that TRIF and IL-1R1 signaling restrain intestinal IgA production locally, through a T-dependent pathway occurring within the PPs.

### TRIF and IL-1R1 shape lymphocyte differentiation within Peyer’s patches

Given the strong correlation between T_FH_ expansion and IgA production^22^, it is important to delineate the cell-intrinsic functions of TRIF or IL-1R1 within lymphocyte populations. To this end, we adoptively transferred splenic naïve CD4 T cells isolated from WT mice into *Trif-*and *Il1r1-*deficient mice. Two weeks later, we analyzed the differentiation status of the donor cells in the PPs (Fig 3A). The activation status of the transferred WT CD4 T cells within the PP were similar regardless of the *Trif* or *Il1r1* expression in the recipient mice (Fig S5A and S5B). Furthermore, we observed that adoptively transferred WT CD4 T cells had a significant increase in T_FH_ differentiation with the PPs in *Trif-* or *Il1r1*-deficient mice compared to the WT transferred cells from the PPs of WT mice (Fig 3B and 3C). This suggests that the T_FH_ differentiation is T cell-extrinsic and is instead driven by the PP microenvironment.

**Figure 3.**
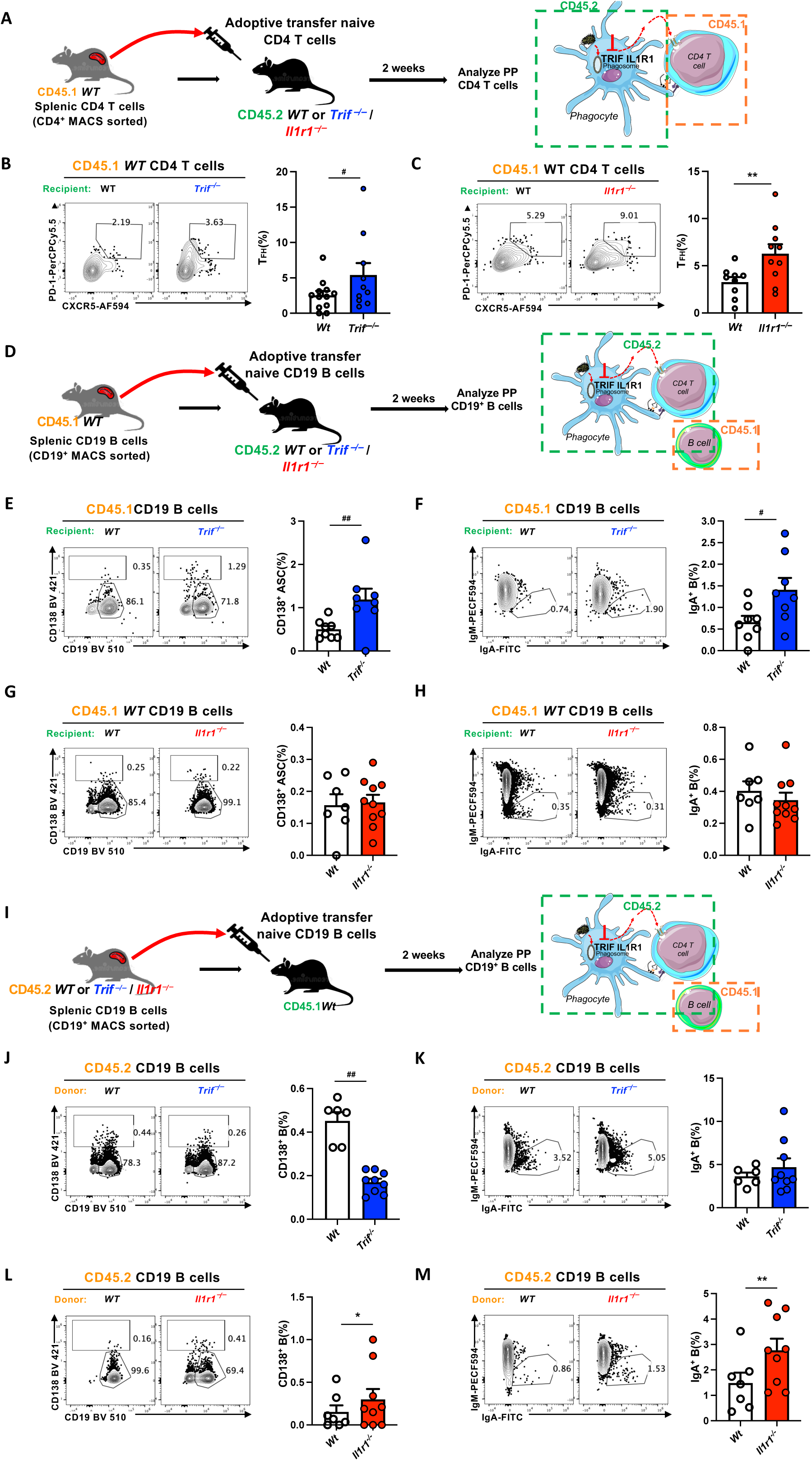
TRIF and IL-1R1 regulate T_FH_ and B cell differentiation in Peyer’s patches. (**A–C**) Adoptive transfer of CD45.1⁺ Wt CD4 T cells into CD45.2⁺ Wt, *Trif^⎻/⎻^*(B), or *Il1r1^⎻/⎻^* (C) recipients. (A) Experimental schematic. (B and C) Representative dot plots (left) and frequencies (right) of donor-derived PD-1⁺CXCR5⁺ T_FH_ cells in Peyer’s patches (PP) two weeks post-transfer. (**D–H**) Wt CD45.1⁺CD19⁺ B cells were transferred into CD45.2⁺ Wt, *Trif^⎻/⎻^*(E, F), or *Il1r1^⎻/⎻^* (G, H) recipients. (D) Schematic. (E and G) Percentages of donor-derived CD138⁺ ASCs in PP. (F and H) Percentages of donor-derived IgA⁺ B cells. (**I–M**) Reciprocal adoptive transfer of CD45.2⁺ Wt, *Trif^⎻/⎻^*(J, K), or *Il1r1^⎻/⎻^* (L, M) CD19⁺ B cells into CD45.1⁺ Wt recipients. (I) Schematic. (J and L) Frequencies of donor-derived CD138⁺ ASCs. (K and M) Frequencies of donor-derived IgA⁺ B cells. Statistical analysis: unpaired Mann–Whitney *U* test;* p<0.05; ** p<0.01; *** p<0.001. *: Wt to *Il1r1^⎻/⎻^* , #: Wt to *Trif^⎻/⎻^* . Each dot represents one mouse. (n = 4-9 per group pooled from two independent experiments)

To further elucidate the cell-intrinsic role of *Trif* and *Il1r1* gene in B cells, we performed reciprocal adoptive transfer experiments using isolated CD19^+^ B cells. We first assessed if the *Trif*- or a *Il1r1*-deficient microenvironment could drive WT B cells toward IgA within PPs. Splenic CD19^+^ B cells from WT mice were transferred into WT , *Trif*- or *Il1r1*-recipients, and donor-derived cells in PPs were analyzed two weeks later (Fig. 3D). The frequencies of ASCs and IgA^+^ B cells among donor-derived cells were comparable between WT and *Il1r1⁻/⁻* recipients (Fig. 3G, H), whereas both B cell subsets were increased when WT B cells were transferred into *Trif⁻/⁻* recipients (Fig. 3E, F). Conversely, to test whether *Il1r1* deficiency confers a B cell–intrinsic advantage, we transferred splenic CD19^+^ B cells from WT or *Il1r1⁻/⁻* donors into WT recipient mice (Fig. 3I). Donor-derived *Il1r1⁻/⁻* B cells exhibited higher frequencies of ASCs and IgA^+^ B cells than WT donor cells (Fig. 3L, M). In contrast, donor-derived *Trif⁻/⁻* B cells did not show an increase compared with WT donor cells (Fig. 3J, K). Of note, the frequencies of ASCs and IgA^+^ B cells among recipient-derived cells were consistent with our prior observations (Fig. S6).

Taken together, these data indicate that IL-1R1—but not TRIF—imposes intrinsic restriction on B cell differentiation toward IgA^+^ and plasma cell fates. These findings support a dual-layered regulatory model in PPs where TRIF signaling shapes the local microenvironment to constrain IgA^+^ B cell expansion (while also limiting T_FH_ expansion), whereas IL-1R1 signaling acts primarily as a B cell–intrinsic brake that raises the activation threshold for class-switch recombination and plasma cell differentiation, thereby preventing excessive IgA responses.

### *Trif*^−*/*−^ and *Il1r1*^−*/*−^ B cells in PP harbor elevated *Ccnd3* expression

To further explore the roles of TRIF and IL-1R1 in T-dependent responses within the PP, we performed single-cell RNA sequencing (scRNA-seq) to profile the transcriptomic landscape for all PP cells. We sorted all live PP cells from *Trif-* and *Il1r1-*deficient mice and from their littermate-control mice (4 mice per group). After analysis by Uniform Manifold Approximation and Projection (UMAP), we identified 19 clusters that were broadly annotated into seven major cell populations (Figs 4A and S7A). These included DCs, naïve B cells, GC IgA⁺ B cells, IgA⁺ plasma cells, CD8 T cells, naïve CD4 T cells, and T_FH_ cells (Fig. 4A). Consistent with our flow cytometry analysis, both KO mice showed the enrichment of T_FH_, IgA producing B cells, and PlasmaBlast/Plasma cells as defined by their transcriptional program compared to their littermate controls (Fig S7A). Gene set enrichment analysis (GSEA) showed the enrichment of an “inflammatory response” signature in WT DCs compared to the *Trif-* and *Il1r1-*deficient counterparts (Fig. S7B). Gene-wise across both *Trif* (y-axis) *and Il1r1* (x-axis) deficiencies we identified several genes differentially expressed in both KO strains (Fig. 4B). Notably, *Ccnd3,* which isan important gene for GC formation^49,50^, was significantly upregulated in several B cell subsets (Fig 4C, Fig. S7C, D), including naïve B cell clusters of both KO mice comprared to their littermate controls. On the other hand, *Fcmr*— a known negative regulator of GC responses, was significantly downregulated^51^ (Fig. S7E). Together, these findings suggest that PP B cells, including naïve B cells, are transcriptionally poised toward GC differentiation.

**Figure 4.**
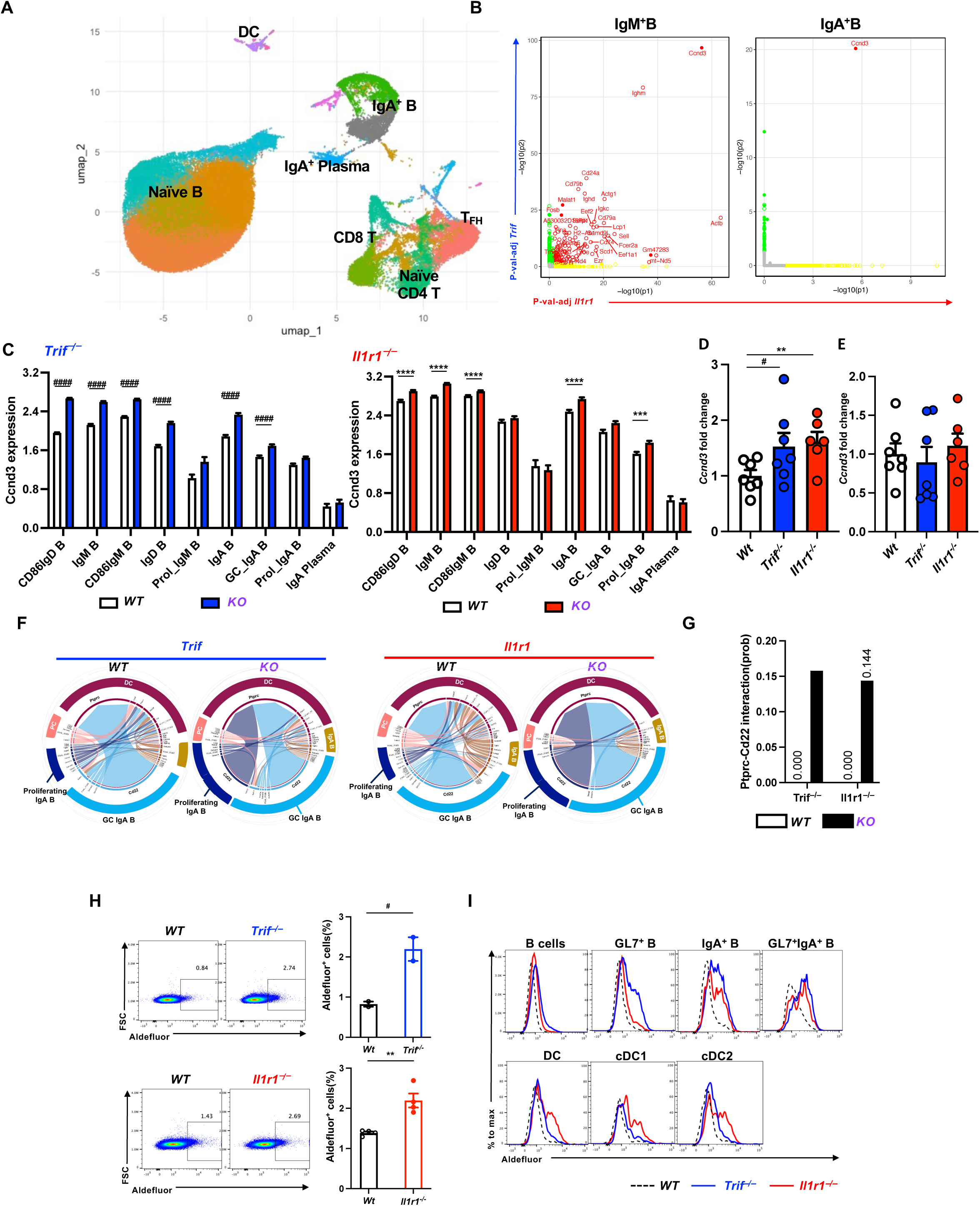
B cells from *Trif^−/−^* and *Il1r1^−/−^* mice are poised to become GC IgA B cells in Peyer’s patches. (**A**) Uniform manifold approximation and projection (UMAP) plot displaying 105,732 cells colored by shared nearest neighbor clusters collected. (**B**) Representative scatter plot comparing gene-wise −log₁₀(adjusted p) values in *Trif* (y-axis) and *Il1r1* (x-axis) comparisons of IgM^+^ B and IgA^+^ B cluster. (**C**) Expression of *Ccnd3* mRNA bar plot in the indicated clusters between *Trif^⎻/⎻^* (left)/*Il1r1^⎻/⎻^*(right) mice and their Wt littermate control. (**D-E**) *Ccnd3* expression in sorted PP (D) and SPL (E) IgM/D B cells measured by qPCR. (**F**) CellChat-derived chord diagrams showing ligand–receptor interactions between IgA^+^ B cells and dendritic cells (DC) in *Trif^⎻/⎻^* (left)/*Il1r1^⎻/⎻^*(right) mice compared to controls. (**G**) Interaction probabilities between PTPRC-CD22 pairs shown in (F). (**H**) Dot plots (left) and percentages (right) of ALDH^high^ cells in *Trif^⎯/⎯^* (top panel) and *Il1r1^⎯/⎯^* (bottom panel) mice compared to wild-type (Wt) controls. (n=2-4 per group) (**I**) Aldefluor histograms of *Trif^⎯/⎯^* and *Il1r1^⎯/⎯^* mice compared to Wt, separated by DC and B cell subtype. Statistical analysis: unpaired Mann–Whitney *U* test; * p<0.05; ** p<0.01; *** p<0.001.

To further validate the *Ccnd3* expression pattern observed in the scRNA-seq analysis, we examined *Ccnd3* expression in IgM⁺ IgD^+^ B cells sorted from the PP of WT and both KO mice. Quantitative reverse transcription PCR (RT-PCR) confirmed that *Ccnd3* was specifically upregulated in PP-derived IgM⁺ B cells from both KO mice (Fig. 4D), while its expression remained similar in splenic B cells from the three groups of mice (Fig. 4E), suggesting that *Ccnd3* upregulation in naïve B cells is restricted to PPs of KO mice.

Beyond cell-intrinsic gene expression, we utilized CellChat, a ligand-receptor interaction analysis tool to assess the PP cell interaction within KO and WT microenvironments. Strikingly, both *Trif⁻/⁻* and *Il1r1⁻/⁻* cells displayed elevated B cell–DC interactions mediated through the CD45 (*Ptprc*)–CD22 axis (Fig 4F, G) compared to WT control cells, as well as enhanced T_FH_–DC interactions via the ALCAM–CD6 axis (Fig S8A, B). These data highlight that both *Trif* and *Il1r1* deficiencies enhance DC-lymphocyte interactions through similar mechanisms to promote GC formation and IgA production, specifically within the PPs.

We previously reported that TRIF and IL-1R1 promoted T_FH_ differentiation and GC formation in a mouse model of parenteral immunization^42^, we speculate that other factors were contributing to these events specifically within the PPs. Previous studies have shown that gut-associated lymphoid tissues, including PPs, have a uniquely high capacity for local RA production compared with peripheral lymphoid organs studies^52^. In addition, RA signaling has been reported to intersect with *Ccnd3*-associated programs in B cells^53,54^. Based on these findings, we hypothesized that the elevated *Ccnd3* expression in both KO strains may be linked to enhanced RA metabolism within the PP microenvironment. To this end, we evaluated bulk PP mRNA expression of those genes associated with RA metabolism. We found that *Aldh1a1*, *Adh1* and *Adh2*, which can increase the level of RA, were markedly upregulated in *Il1r1-*deficient mice while they were modestly upregulated in *Trif-*deficient mice (Fig. S8D).

Moreover, we observed that PP cells from *Trif-* and *Il1r1-*deficient mice showed higher retinaldehyde dehydrogenase activity (Fig 4H). Further analysis demonstrated that the enzymatic activity increased specifically in some *Trif⁻/⁻* and *Il1r1⁻/⁻* PP cells compared to their WT counterparts, such as within IgA^+^ B cell and DC populations (Fig 4I) but not within T cells (Fig. S8E). Altogether, these data suggest that *Trif* and *Il1r1* deficiencies are associated with enhanced RA production, which may contribute to the increased IgA production observed in the intestines^26,45^. This finding is consistent with our previous observations from T cell adoptive-transfer experiments and with CellChat-inferred B cell–DC interactions.

### *Trif-* and *Il1r1*-deficient mice produce higher level of Cholera-toxin-specific IgA in their intestines after CT vaccination

We next asked whether the absence of TRIF or IL-1R1 signaling helps to elicit stronger mucosal antigen-specific TD IgA immunity within the intestines following exposure to an exogenous antigen. The cholera toxin (CT) vaccination model is a well-characterized model that relies on immune activation in the PPs and the induction of T_FH_-dependent anti-CT IgA^55^. We orally administered CT to WT, *Trif*-, and *Il1r1-*deficient mice, then evaluated the level of stool and serum CT-specific IgA/G at the indicated time points (Fig 5A). Four weeks after the initial vaccination, CT-specific IgA levels in the stool were significantly increased in both *Trif*- and *Il1r1*-deficient mice by more than threefold (Fig. 5B, Fig. S9A). However, serum CT-specific IgG levels remained unchanged (Fig. 5C, Fig. S9B). Serum CT-specific IgA and stool CT-specific IgG were below the limit of detection in most samples and were therefore not included in the analysis. Altogether, these findings indicate that enhanced intestinal IgA production observed in *Il1r1-* and *Trif-*deficient mice is beyond steady-state, SPF conditions, and it is equally important in the context of mucosal IgA response in the intestines following oral vaccination.

**Figure 5.**
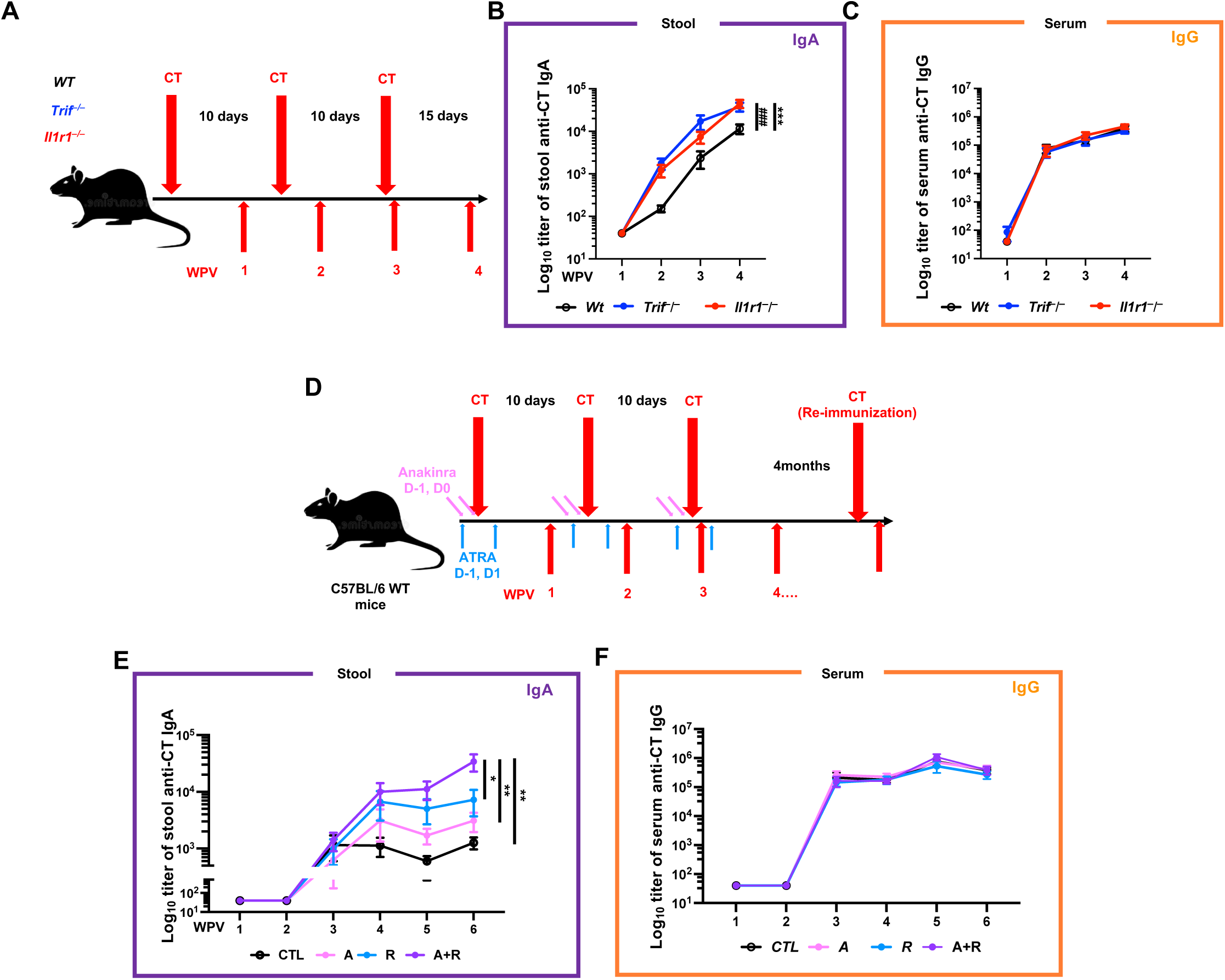
Enhanced antigen-specific IgA responses following cholera toxin vaccination in absence or blockade of the TRIF/IL1R1 pathway. **A**) Schematic representation of the vaccine delivery (n = 12-16 per group pooled from three independent experiments). (**B**) Stool CT-specific IgA levels over time following the first vaccination. (**C**) Serum CT-specific IgG levels over time following the first vaccination. (**D**) Schematic representation of the vaccination protocol. The strategy for administering anakinra and all-trans retinoic acid (ATRA) is described in the Methods section (n = 10-12 per group pooled from two independent experiments). (**E**) Stool CT-specific IgA levels over time following the first vaccination. (**F**) Serum CT-specific IgG levels over time following the first vaccination. CT: cholera toxin; CTL: control; A: anakinra; R: ATRA; A+R: anakinra plus ATRA; WPV: weeks post the first vaccination. Statistical analysis was performed using two-way ANOVA. *p < 0.05; **p < 0.01; *** p < 0.001. *: *Wt* to *Il1r1*^⎯^*^/^*^⎯^, *#*: *Wt* to *Trif^⎯/⎯^* .

Collectively, our results demonstrate that genetic disruption of the TRIF/IL-1R1 pathway selectively enhances mucosal IgA responses while leaving systemic CT-specific IgG levels unchanged. To explore the translational potential of these findings, we investigated whether pharmacological inhibition of IL-1R1 could recapitulate phenotypes observed in the genetically-engineered mice. Given the evidence implicating RA metabolism, and the established role of RA in promoting intestinal IgA responses^25,56^, we decided to combine Anakinra, a clinical-grade IL-1R1 antagonist, with all-trans retinoic acid (ATRA) to see if we could augment mucosal IgA immunity. Treatment with ATRA or Anakinra and ATRA significantly increased intestinal antigen-specific IgA titers without enhancing serum IgG levels (Figs. 5D–F and S9D, E). The combination treatment of Anakinra and ATRA produced the strongest mucosal IgA response and significantly increased IgA titers compared with both the control and ATRA-alone groups. At the peak of the response, we observed up to 66 times more antigen-specific IgA antibodies in the stool of mice treated by the combination of Anakinra and ATRA 6 weeks post vaccination compared to the untreated vaccinated controls. Notably, stool CT-specific IgA titers did not correlate with serum CT-specific IgG titers (Fig. S9C), indicating that variation in mucosal IgA responses is uncoupled from the magnitude of the systemic humoral response. Furthermore, this enhanced mucosal response was sustained even after re-immunization four months after primary vaccination (Figs. S9F, G). Altogether, these findings underscore a targetable pathway to augment mucosal immunity.

## DISCUSSION

Innate inflammatory pathways are classically viewed as positive regulators of adaptive immunity and are widely exploited in vaccine adjuvant design^57^. Unexpectedly, our findings demonstrate that TRIF- and IL-1R1-associated pathways instead function as restraining checkpoints for intestinal IgA production under both steady-state and vaccination conditions. These observations reveal a fundamental divergence between systemic and mucosal humoral regulation and suggest that inflammatory signaling can impose tissue-specific constraints on IgA differentiation within the intestinal immune niche. Importantly, pharmacological modulation through combined administration of Anakinra and ATRA markedly increased antigen-specific IgA in WT mice after CT immunization. Together, these findings define an innate immune checkpoint that limits the magnitude of mucosal IgA responses and provide proof-of-concept that transient IL-1R1 inhibition in combination with RA supplementation, can be leveraged to enhance oral vaccine–induced IgA immunity (Fig. S10).

Mucosal IgA is increasingly recognized as a correlate of protection against pathogens entering through respiratory or intestinal surfaces, yet most licensed vaccines are delivered intramuscularly or subcutaneously, which reliably induces systemic IgG but comparatively limited mucosal IgA^58,59^. A major advance in mucosal vaccinology has been the development of “prime-and-pull” strategies, which have demonstrated that local tissue cues can be harnessed to establish potent site-specific immunity^36–38^. These approaches have been particularly influential in showing that systemic priming followed by local recruitment signals can generate robust tissue-resident memory T cell (T_RM_) responses at mucosal surfaces. Building on this concept, our findings identify the TRIF/IL-1R1 regulatory axis as an endogenous checkpoint that constrains mucosal IgA responses during oral immunization. Our data suggest that modulation of this axis can robustly amplify antigen-specific intestinal IgA responses directly through oral immunization. Thus, our work extends the principle of local immune programming to the humoral arm of mucosal immunity and suggests that targeting innate checkpoints may provide a complementary approach for enhancing oral vaccine efficacy. Since prime-and-pull strategies have primarily been optimized for long-term mucosal immunity, particularly to “pulling” T_RM_ within mucosal niches, our study highlights an additional opportunity to tune local B cell differentiation and enhance mucosal IgA production. A remaining question is whether TRIF/IL-1R1 signaling similarly shapes antigen-specific T_RM_ formation within the same niches. Defining how this axis coordinates mucosal memory niches is an important direction for future work.

Prior work has proposed that the TRIF/IL-1R1 axis contributes to sensing “viability-associated” cues during systemic vaccination^42^, and viability-related signals have also been implicated in promoting intestinal IgA responses^60^. We initially hypothesized that this axis would similarly promote mucosal IgA responses. However, our genetic and pharmacological data instead support a model in which TRIF/IL-1R1 signaling acts as a localized restraint on PP-driven IgA differentiation. This finding highlights a context-dependent role for IL-1 signaling in shaping humoral immunity. Recent work showed that an AIN-93G purified diet markedly reduces PP T_FH_ cells and downstream IgA responses in post-weaning mice, whereas soy supplementation restores this axis through IL-1β signaling^61^. Together with our findings, these observations support a model in which IL-1 signaling functions as a context-dependent molecular switch rather than a uniformly pro-IgA pathway. Depending on dietary and microbial inputs, tissue niche, and the cellular target of IL-1R1 signaling, this pathway may either promote or restrain Tfh/GC-associated humoral responses. Defining the conditions under which TRIF-and IL-1R1-associated signaling supports versus constrains PP immunity will be important for understanding how innate cues tune mucosal IgA responses. A critical next step will be to determine how this ’brake’ is engaged within the unique mucosal landscape—specifically, whether distinct microbial contexts, such as defined colonization or germ-free conditions, modulate the requirement for TRIF/IL-1R1 in shaping IgA magnitude and repertoire specificity. Notably, both human and mouse B cells are known to express IL-1R1, particularly after activation^62,63^. However, the biological function of B cell IL-1R1 has remained poorly defined. Our data supports a model in which IL-1R1 directly restrains IgA differentiation through a B cell-intrinsic mechanism.

Mechanistically, our single-cell analyses highlight candidate programs and cellular interfaces that may contribute to enhanced IgA output under TRIF/IL-1R1 deficiency. At the B cell level, we observed downregulation of *Fcmr*^51^, which has been reported to restrain GC responses, together with enrichment of cell-cycle–associated programs, including *Ccnd3*, a gene linked to GC B cell proliferation^49,50,64,65^. Together, these transcriptional changes suggest that loss of either *Trif* or *Il1r1* shifts PP B cells toward a state more permissive for GC entry and expansion. In parallel, CellChat inference suggested increased B cell–DC communication through the CD45–CD22 axis in *Trif-* and *Il1r1*-deficient conditions relative to WT (Fig. 4 F). CD22 (Siglec-2) tunes BCR signaling thresholds and proliferative responses^66^, but whether CD45 presented *in trans* by DCs functionally modulates CD22-dependent signaling or B cell fate decisions remains to be established and will require orthogonal experimental validation.

Within the PPs, blocking IL-1R1 transiently relieves an inhibitory checkpoint on GC-associated B cell differentiation, allowing RA-associated signals to more effectively support IgA production. This synergy is clinically relevant because IL-1R1 inhibition might lower the required dose of retinoids, potentially mitigating the toxicity issues often seen with systemic RA treatment^28^. In our current experiments, Anakinra was administered intraperitoneally; future studies should determine whether mucosa-targeted delivery, such as oral or intranasal formulations, can enhance local efficacy while minimizing systemic exposure. Finally, meta-analyses of rheumatoid arthritis trials suggest that Anakinra does not significantly increase the risk of serious infections compared with the placebo, although this endpoint does not capture all aspects of mucosal defense^67^, and IL-1R blockade has shown context-dependent effects on mucosal injury and inflammation^68,69^. Determining how IL-1R1 antagonism reshapes mucosal antibody quality, including IgA repertoire features, as well as IgA–microbiota interactions^43^, will be important for translating this strategy for mucosal vaccinology.

### Limitations of the study

This study has some limitations. First, all *in vivo* experiments were performed in mice, and the effects of TRIF/IL-1R1 modulation may differ in humans. Although Anakinra and ATRA are clinically prescribed drugs that target conserved inflammatory and RA-associated pathways, their effects on human mucosal vaccine responses remains unknown. In particular, dosing, route of administration, safety, and the balance between systemic and local mucosal effects will need to be carefully evaluated before this strategy can be translated to humans. Thus, our pharmacological data should be interpreted as proof-of-concept evidence that IL-1R1 blockade and RA-associated signaling can enhance oral vaccine-induced IgA responses in mice.

Second, while our genetic and adoptive transfer data identify IL-1R1 as a B cell–intrinsic brake, the exact cellular source of TRIF-dependent regulation remains to be fully elucidated. Our findings suggest that TRIF acts extrinsically to B and T cells to shape the PP microenvironment. Identifying the specific TRIF-expressing compartment—whether antigen-presenting cells, stromal cells, or epithelial-associated cells—will require extensive further investigation using a broad panel of cell-type-specific *Ticam1*-deficient models. Such work will be an important next step in defining how the innate microenvironment coordinates GC-associated IgA responses.

## RESOURCE AVAILABILITY

### Lead contact

Requests for further information should be directed to the lead contact, Gaetan Barbet

### Materials availability

This study did not generate new reagents.

### Data and code availability

Mouse single-cell RNA is deposited in NCBI’s Gene Expression Omnibus 90 and are accessible through the GEO Series accession number GEO: GSE328882. This study did not generate any new code.

## ACKNOWLEDGMENTS

We thank professor Emilie K. Grasset for providing *Taci^−/−^* mouse strains. This work was supported, in part, by research grants from Robert Wood Johnson Foundation Grant Project (RWJF Grant Project # 826399 to G.B.) and the Basic research award from the Feldstein Medical Foundation; Moderna Global Fellowship 2022 (DR#20892 to C.C.S.); and the National Institutes of Health/National Heart, Lung, and Blood Institute (R01HL164404 to S.S.A). G.B. was supported by the Career Development Award (#602723) from the Crohn’s and Colitis Foundation. The laboratory thanks the Robert Wood Johnson Foundation Grant #74260 for their support of the CHINJ.

## AUTHOR CONTRIBUTIONS

Conceptualization, C.C.S. and G.B.; methodology, C.C.S. and G.B.; investigation, C.C.S., N.G.B., M.S., C.J, and N.G.; visualization, C.C.S., C.T., and G.B.; formal analysis, C.C.S., C.T., S.H.C., and Q.Y.; funding acquisition, C.C.S., S.S.A and G.B.; supervision, A.R., and G.B.; writing – original draft, C.C.S.; writing – review and editing: all authors.

## DECLARATION OF AI AND AI-ASSISTED TECHNOLOGIES

During the preparation of this work, the authors used ChatGPT in order to improve the language and grammar of the manuscript. After using this tool, the authors reviewed and edited the content as needed and take full responsibility for the content in this published article.

## EXPERIMENTAL MODEL AND SUBJECT DETAILS

### Mice

C57BL/6J, *Trif^−/−^*, *Il1r1^−/−^*, *Il1r1^f//fl^*, CD45.1 and *Itgax*-CRE mice were purchased fromJackson Laboratories. *Taci^−/−^*mice was kindly provided by Emilie K. Grasset, Ph.D. (Weill-Cornell Medicine, NY). Unless otherwise mentioned, experiments were done using 8-12 week old mice (age-matched). Both sexes were included to account for possible biological variability. Mice were euthanized by carbon dioxide asphyxiation, followed by cervical dislocation. Mice were in the Child Health Institute of New Jersey (CHINJ) animal facility. Animal care and experimental procedures were carried out following the guidelines of the Institutional Animal Care and Use Committee (IACUS) of Rutgers University and the National Institutes of Health Guide for the Care and Use of Laboratory Animals.

### Fecal IgA Flow Cytometry

Fecal pellets collected directly from indicated mice at indicated time points in 1 ml Phosphate Buffered Saline (PBS) per 100 mg fecal material on ice. Fecal pellets were homogenized and then centrifuged (300 x g, 5 min, 4 °C) twice to remove large particles. Fecal bacteria in the supernatants were centrifuged (8000 x g, 5 min, 4 °C) to spin down. The supernatants were kept at -30 °C for further ELISA analysis. The pellets were resuspened with 1 ml ice-cold PBS. A sample of this bacterial suspension (2 µl) was stained with with 50 µl Magnetic-activated cell sorting (MACS, 2% FBS and 1 mM EDTA in DPBS) staining buffer containing APC-conjugated anti-mouse IgA (1:100; ThermoFisher clone mA-6E1) and syto 9 (1:500; ThermoFisher) for 30 min on ice. Samples were then washed 3 times with 1 ml staining buffer before flow cytometric analysis (LSR, BD).

### Anti-mouse IgA Antibody ELISA

Fecal samples were collected as previously described. Sera were isolated from facial vein blood at the indicated time points. Total IgA and IgG levels were measured by ELISA using 96-well plates coated overnight at 4°C with anti–mouse Ig antibodies. After blocking with 5% BSA in PBS, fecal and serum samples were added and incubated for 2 hours at 4°C. Plates were then washed and incubated with HRP-conjugated rabbit anti–mouse IgA or IgG antibodies (Southern Biotech or Jackson ImmunoResearch) for 1 hour. For cholera toxin (CT)–specific ELISA, plates were first coated with ganglioside GM1 (0.5 nM) overnight at 4°C, followed by incubation with CT (0.5 µg/ml) for 2 hours at room temperature before blocking. Sample incubation and detection were performed as described above. Bound antibodies were visualized using 3,3’,5,5’-tetramethylbenzidine (TMB; Sigma), and absorbance was measured at 450 nm using an Agilent Stratagene microplate reader.

### Lipocalin-2 quantification

The inflammation status of mice was evaluated by measuring the levels of Lipocalin-2 (LCN-2) in fecal supernatants via an ELISA assay (R&D systems, DuoSet ELISA Mouse Lipocalin-2/NGAL). Briefly, fecal samples collected previously were used to perform the LCN-2 ELISA assay following the manufacturer’s instructions. Absorbance was read at 450 nm and the LCN-2 concentration obtained was normalized for the dilution factor and for stool weight.

### Histology

The small intestines of indicated mice were harvested and gently rinsed with modified Bouin’s fixative (a mixture of 50% ethanol and 5% acetic acid in water) using a 10-ml syringe and a gavage needle. Swiss rolls were fixed in 10% buffered formalin overnight at room temperature, then they were rinsed and stored in 70% ethanol at 4 °C. The embedding in paraffin, sectioning (5 µm), and hematoxylin and eosin (H&E) staining of the intestines were done by Nationwide Histology LLC. The histological evaluation was further confirmed by A.B.R. at the Rutgers Child Health Institute of NJ.

### Intestinal permeability

The intestinal permeability assay was performed as described ^70^. Indicated mice were fasted for 2 hours and then gavaged with a mixture of FITC-Dectran 4kd (80 mg/ml; Sigma-Aldrich, 46944). Facial blood was collected in the indicated time points. Sera was collected for the analysis. Fluorescence intensity was determined by using a plate reader at 495-nm excitation / 525-nm emission.

### Quantitative Real Time Reverse Transcription Polymerase Chain Reaction (qRT-PCR)

Total RNA was isolated from total PP cells using Trizol ambion. Contaminating genomic DNA was removed by DNase digestion (DNase I, Roche). RNA (100 ng to 1 µg) was used to synthesize complementary DNA (cDNA) by reverse transcription using the iScript cDNA Synthesis Kit (Bio-Rad). Quantitative PCRs were run with cDNA template in the presence of 0.5 µM forward and reverse primer and 1× solution of Maxima SYBR Green qPCR Master Mix (Thermofisher). A complete list of primer sequences is provided in table S3. All reactions were performed in duplicate and the samples were normalized to β-actin.

### Isolation of leukocytes from LP, MLN and PP

Leukocytes were isolated from the LP, MLNs, and PPs as previously described, with minor modifications. Briefly, PPs and MLNs were collected, and the small intestine, including the duodenum, jejunum, and ileum, was excised. PPs and MLNs were mechanically dissociated using a syringe plunger and filtered through a 100-μm strainer to generate a single-cell suspension in MACS buffer. The small intestine was then opened longitudinally with surgical scissors and flushed with ice-cold PBS to remove fecal contents. Intestinal tissues were cut into 0.5-cm pieces and transferred into 50-ml conical tubes containing 20 ml PBS supplemented with 2% FBS. Samples were vigorously shaken for 10 s using a Vortex-Genie (Scientific Industries) and passed through 250-μm cell strainers. Fresh PBS containing 2% FBS was added, and the shaking and filtering steps were repeated a total of four times. To remove the intestinal epithelial layer, samples were washed three times with 20 ml of prewarmed MACS buffer and passed through 250-μm cell strainers each time. Samples were then washed with ice-cold PBS containing 2% FBS to remove residual EDTA from the MACS buffer. Tissues were subsequently resuspended in RPMI 1640 containing 10% FBS, 0.5 mg/ml collagenase D (Roche), and 1 mg/ml DNase I (Roche), and incubated at 37 °C with shaking for 30 minutes. After digestion, samples were passed through an 18-gauge needle and then filtered through 100-μm nylon cell strainers (BD Falcon) to obtain a single-cell suspension. Leukocytes were enriched using a 37.5% Percoll (Sigma) density gradient according to the manufacturer’s instructions. Following centrifugation, leukocytes were resuspended in MACS buffer for flow cytometric analysis.

### Flow cytometric analyses

All samples were pretreated with Zombie NIR (Biolegend) for 10 min at 4 °C to discriminate viable cells. Next, samples were treated with Fc block for 10 min at 4 °C followed by fluorescently-conjugated antibody labelling at 4 °C for 60 min. The following antibodies were used for these studies: The antibodies include fluorochrome-conjugated monoclonal antibodies to mouse IgM (R6-60.2), Bcl-6 (K112-91), CD21/CD35 (7G6), PSGL1 (2PH1)(all from BD Bioscience); CD3 (17A2), CD4 (RM4-5), CD4 (RM4-5), CD8 (53-6.7), CD11b (M1/70), CD24 (M1/69), CD44 (IM7), CD45 (30-F11), CD45 (30-F11), TACI (8F10), MHCII (30-F19), NK1.1 (PK136), CD45.2 (104), CD19 (6D5), CD11c (N418), CD103 (20000000), PD-1 (29F.1A12), CD138 (281-2), GL7 (GL7), CXCR5 (2G8), CX3CR1 (SA011F11), ICOS (C398.4A), CD23 (B3B4), Tbet (4B10), GATA3 (16E10A21), IgA (MA-6E1), IgA (MA-6E1)(all from Biolegend); FOXP3 (FJK-16s), CD11c (N418), CD25 (PC61.5), CD45.1 (A20), MHC Class II (I-A/I-E) (M5/114.15.2), RORγT (820) (all from Invitrogen). Raldh activity was determined using the Aldefluor® assay (StemCell technology, Vancouver, Canada), as described. Briefly, cells suspension (1×10^6^ cells/ml) were incubated for 45 minutes at 37°C in Aldefluor assay buffer containing activated Aldefluor substrate in the presence or absence of Raldh inhibitor diethylaminobenzaldehyde (DEAB). The cells were subsequently stained with specific antibodies, washed, resuspended in Aldefluor assay buffer and analyzed using a Cytek Aurora flow cytometer(Cytek).

### Adoptive Cell Transfer

Reciprocal adoptive transfer experiments were performed to assess the contributions of T and B cells. For T cell transfer, CD4^+^ splenocytes were purified by magnetic separation according to the manufacturer’s instructions (Miltenyi Biotec). In one setting, CD4^+^ T cells from congenically marked C57BL/6 CD45.1 mice were intravenously transferred into C57BL/6 CD45.2, *Trif^−/−^* or *Il1r1^−/−^*recipient mice (9 × 10^6^ cells per mouse). In the reciprocal setting, CD4^+^ T cells from C57BL/6 CD45.2, *Trif^−/−^* or *Il1r1^−/−^* mice were transferred into C57BL/6 CD45.1 recipient mice (9 × 10^6^ cells per mouse). For B cell transfer, CD19^+^ splenocytes were purified by magnetic separation according to the manufacturer’s instructions (Miltenyi Biotec). In one setting, CD19^+^ B cells from C57BL/6 CD45.1 mice were intravenously transferred into C57BL/6 CD45.2, *Trif^−/−^* or *Il1r1^−/−^*recipient mice (9 × 10^6^ cells per mouse). In the reciprocal setting, CD19^+^ B cells from C57BL/6 CD45.2, *Trif^−/−^* or *Il1r1^−/−^* mice were transferred into C57BL/6 CD45.1 recipient mice (9 × 10^6^ cells per mouse). Recipient mice were analyzed 2 weeks later.

### Single-cell RNA sequencing

#### Nuclei isolation and fixation

Live cells were isolated from the PPs of *Trif-* and *Il1r1-*deficient mice and their littermate WT controls, and then sorted using a Cytek Aurora CS cell sorter. Each group consisted of four mice, including two males and two females housed in different cages. 1 x 10^6^ cells from each sample were fixed with ScaleBio Sample Fixation Kit. The fixed samples were stored at −80 °C until all samples were collected.

### Single-nucleus RNAseq library preparation and sequencing

Frozen fixed cells were thawed on ice and counted twice via hemocytometer. Using the ScaleBio Single Cell RNA Sequencing Kit v1.1 (Scale Biosciences), all 16 samples were prepared in one batch. Cells were loaded at 10,000 per well (6 wells/sample) to add RT barcodes as sample identifiers. Pooled cells were distributed across the 384-well plate for Ligation Barcode addition, then pooled again with 1600 nuclei per well for second strand synthesis. After tagmentation and indexing PCR, libraries were pooled (5μl each), cleaned with 0.8X SPRIselect beads (Beckman Coulter B23317), and fragment size/concentration quantified using High Sensitivity D5000 Screentape (Agilent). Sequencing was performed on a NovaSeq X Plus (Illumina) by Admera Health.

### Single-nucleus RNAseq data processing

Fastq files were demultiplexed, combined, and processed using ScaleBio’s nextflow pipeline. Filtered count matrices were imported using Seurat v4.3.042 to create Seurat objects for downstream analyses. The final dataset contained 105,732 cells. PCA was performed. Dimensionality reduction used UMAP. Clustering was performed using the Louvain algorithm (resolution 0.01-2.4), with optimal resolution (2.0) determined by clustrer v0.5.044.

### Immunizations

Mice were randomized to treatment groups before first treatment using block randomization based on body weight. Sample collection was performed at a consistent time of day, and ELISA quantification was conducted with the operator blinded to group identity. Each group was housed in two independent cages, and cage-level inspection indicated that the observed group differences were not driven by a single cage. Mice were orally immunized with 10 µg of cholera toxin (CT, Sigma) dissolved in 0.1 ml of phosphate-buffered saline (PBS) containing 3% (w/v) NaHCO_3_. Three oral immunizations were given, as indicated, with 10 days between immunizations. Analysis of immune responses was done 4 weeks after the final immunization. For the Anakinra experiment, Anakinra and Anakinra plus ATRA group mice received Anakinra (1 mg per mouse; MCE^40^) twice intraperitoneally on D-1 before immunization to D-0 after immunization. ATRA and Anakinra plus ATRA group mice orally received 300 µg ATRA per mouse^71^ in soybean oil/mouse on D-1 before immunization and D-1 after immunization.

### Statistical analysis

Statistical analyses were performed as indicated in the figure legends using GraphPad Prism (version 11.0.0; GraphPad Software). Two-tailed unpaired Student’s t test or Mann–Whitney U test was used as appropriate. Data are presented as mean ± s.e.m. Significance levels: *P < 0.05, **P < 0.01, ***P < 0.001.

## STAR★METHODS

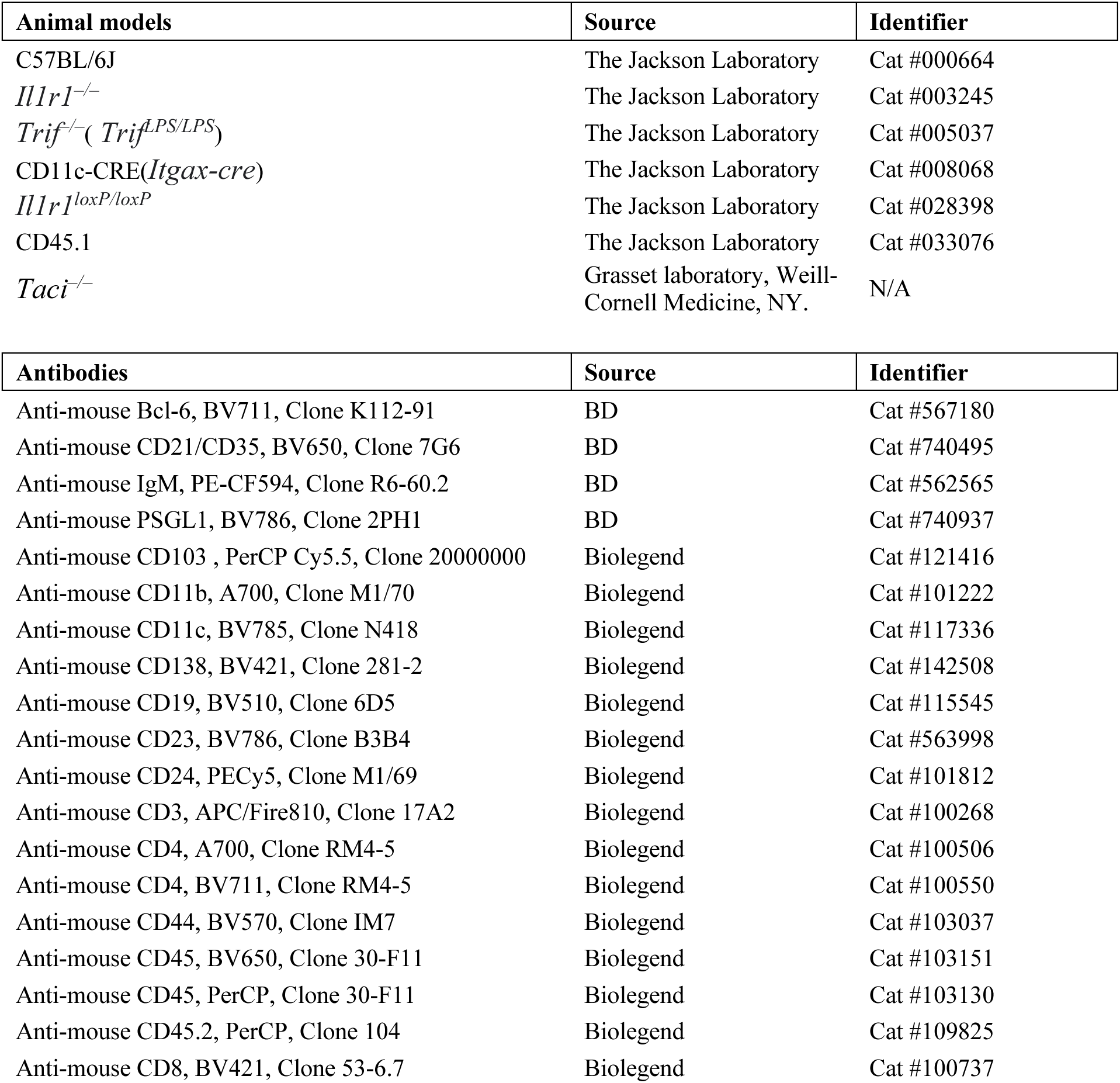
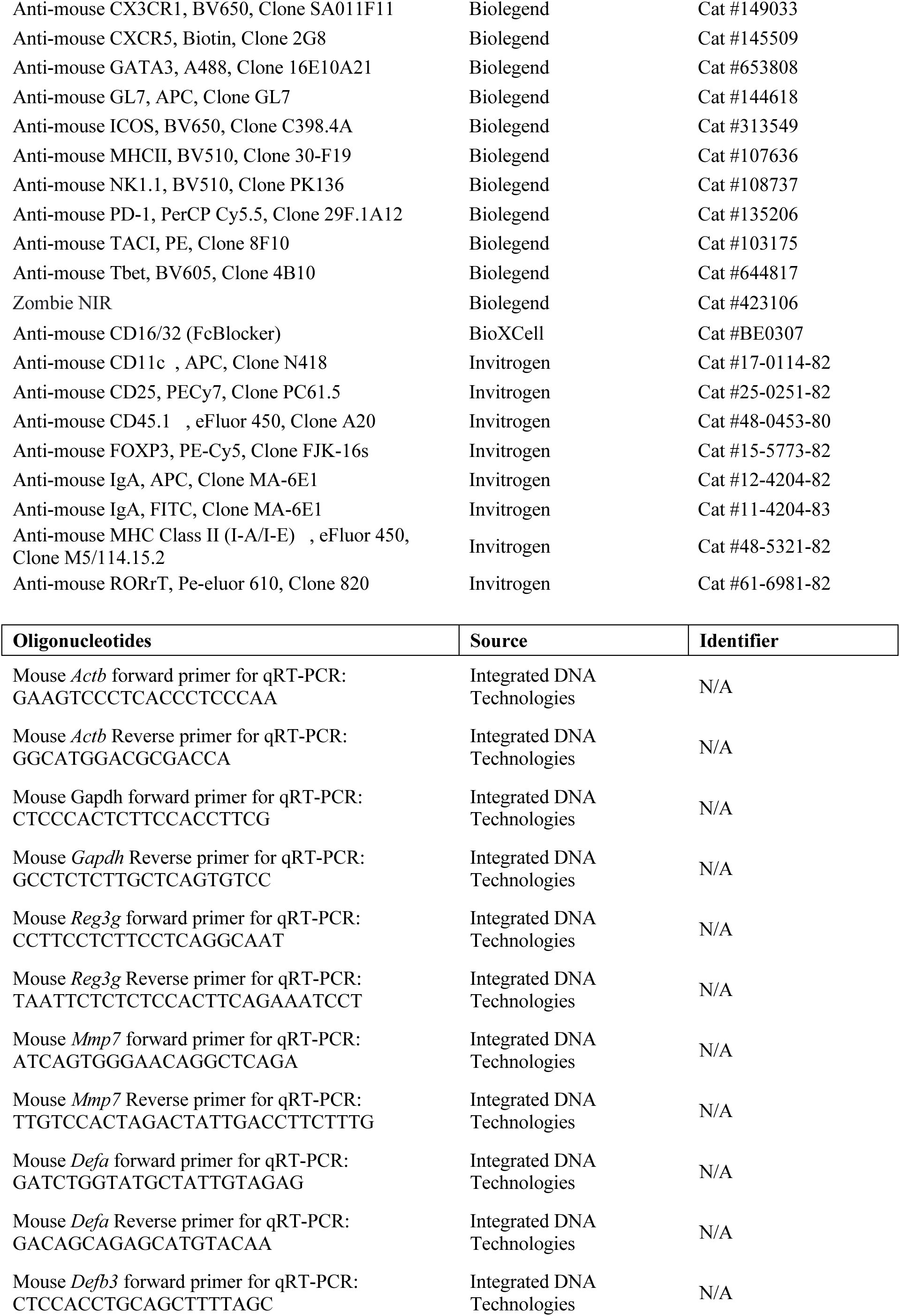
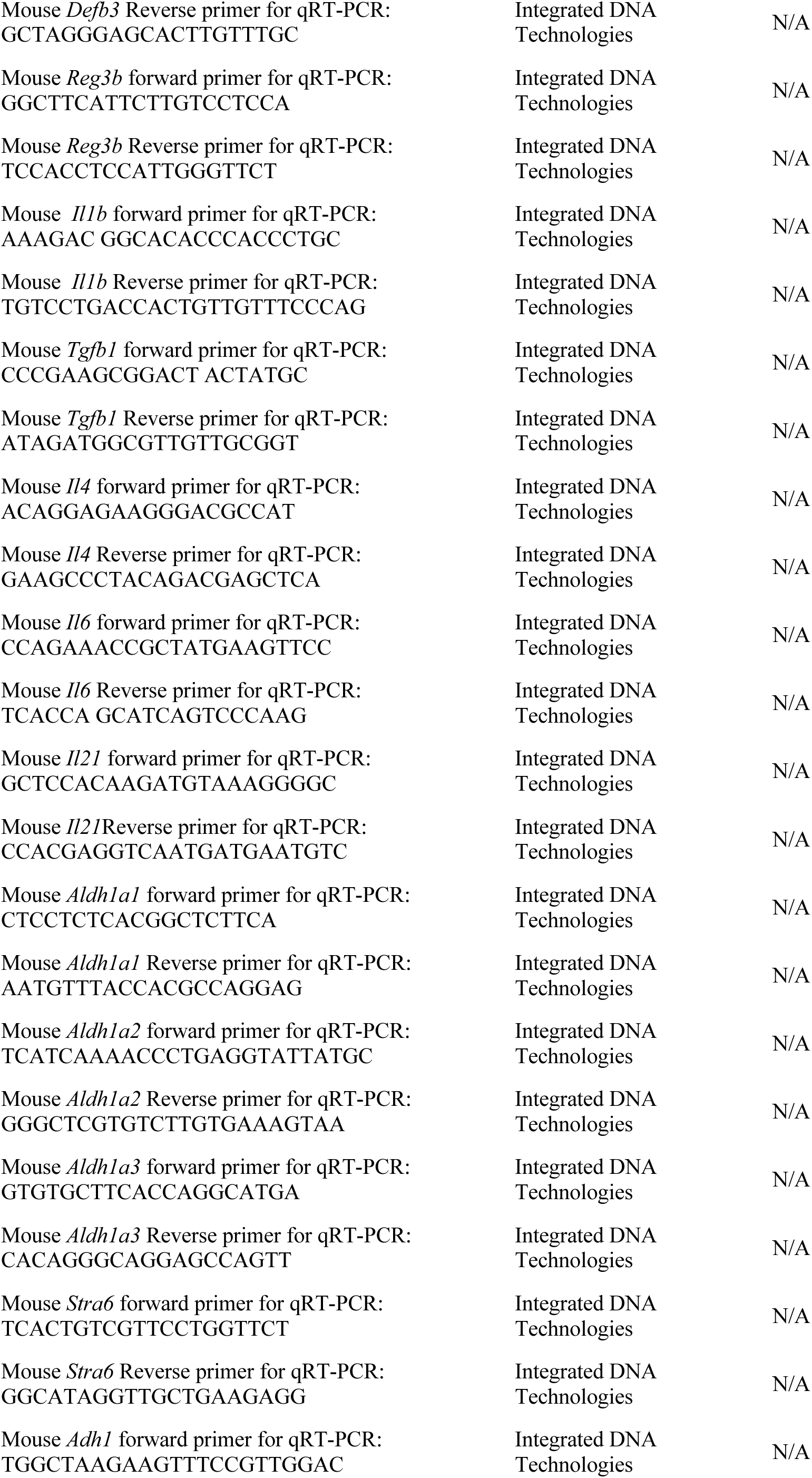
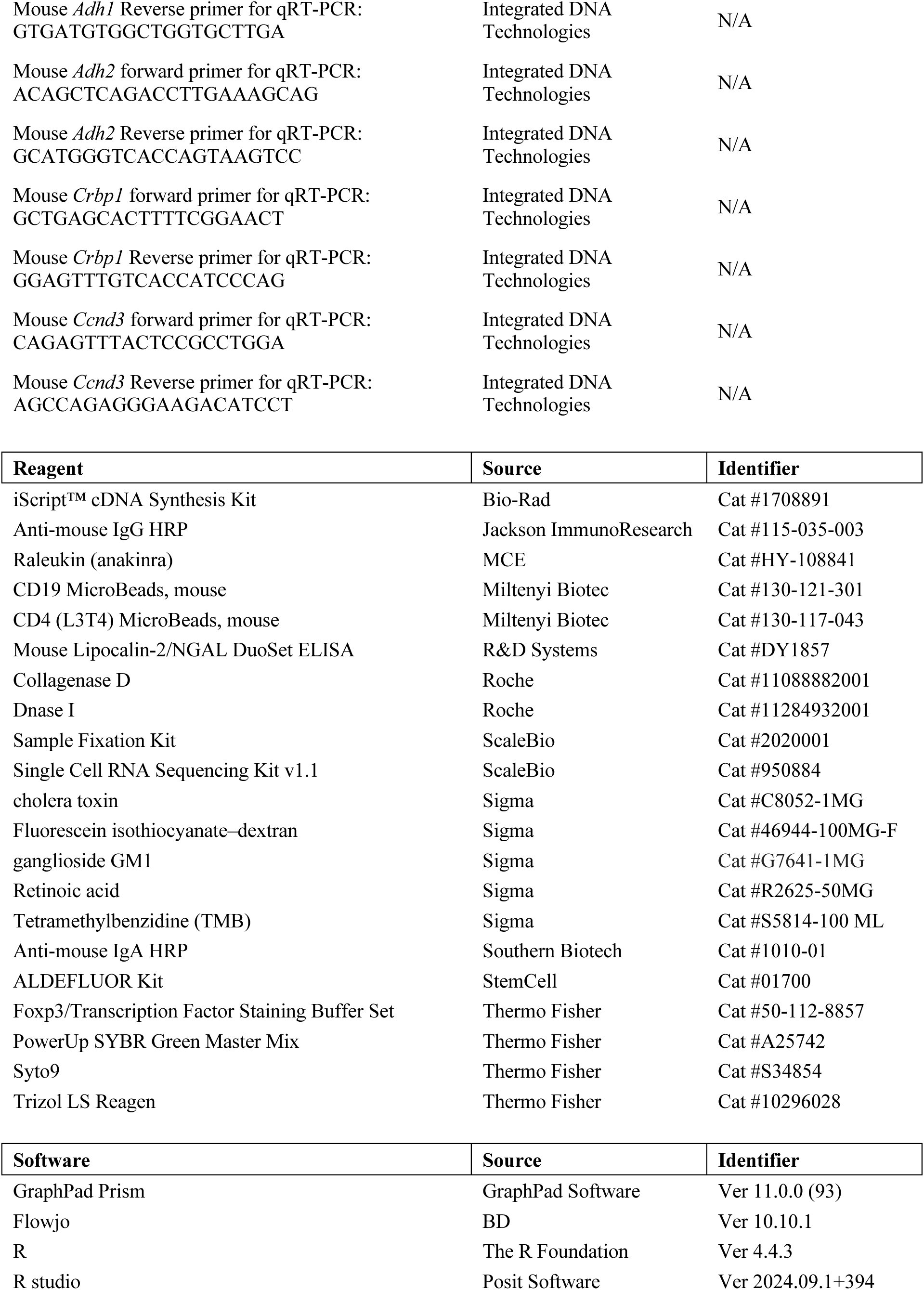
KEY RESOURCES TABLE.

