## Supplemental figures for "A regulatory TRIF/IL-1R1 axis controls T-dependent IgA production in the intestines"

### Figure S1

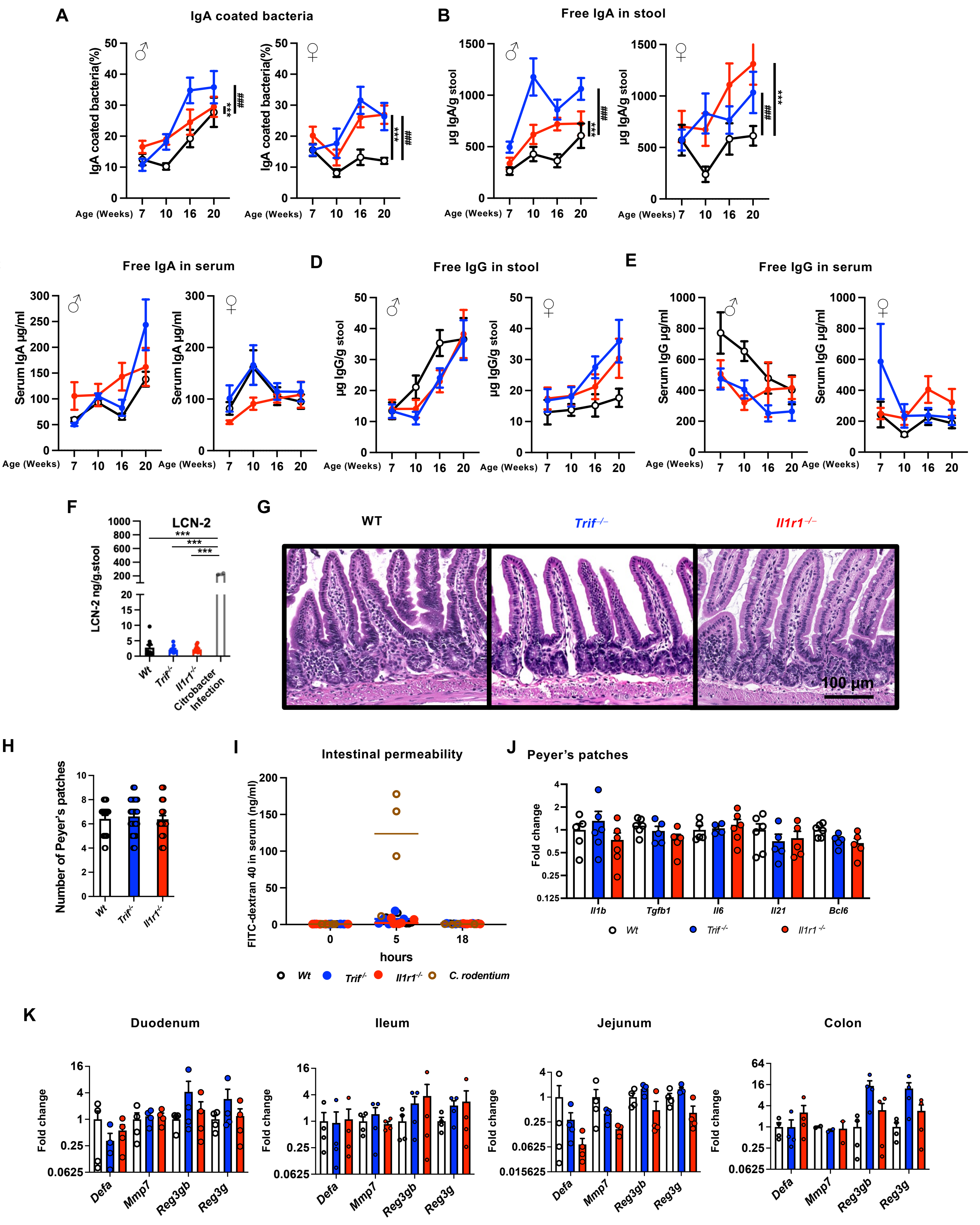
A
IgA coated bacteria
IgA coated bacteria(%)
Age (Weeks)
B
Free IgA in stool
µg IgA/g stool
C
Free IgA in serum
Serum IgA µg/ml
D
Free IgG in stool
µg IgG/g stool
E
Free IgG in serum
Serum IgG µg/ml
F
LCN-2
LCN-2 ng/g.stool
Wt
Trif-/-
Il1r1-/-
Citrobacter Infection
G
WT
Trif-/-
Il1r1-/-
100 µm
H
Number of Peyer's patches
I
Intestinal permeability
FITC-dextran 40 in serum (ng/ml)
hours
C. rodentium
J
Peyer's patches
Fold change
Il1b
Tgfb1
Il6
Il21
Bcl6
K
Duodenum
Ileum
Jejunum
Colon
Defa
Mmp7
Reg3gb
Reg3g

**Supplementary Figure 1. Absence of intestinal inflammation in *Trif*- and *Il1r1*-deficient mice under steady-state conditions.**

(**A**) Percentage of IgA-coated bacteria over time (WT♂=16, ♀=13; *Trif*$^{-/-}$ ♂=13, ♀=12; *Il1r1*$^{-/-}$♂=15, ♀=13 pooled from four experiments). (**B, C**) Free IgA levels in stool (B) and serum (C), measured by ELISA. (**D, E**) Free IgG levels in stool (D) and serum (E), measured by ELISA. (**F**) Stool lipocalin-2 (LCN-2) levels in indicated mice (n = 5 per group). Wild-type mice infected with *C. rodentium* for 9 days served as positive controls. (**G**) Hematoxylin and eosin (H&E) staining of ileal sections from 10-week-old mice. (**H**) Number of Peyer's patches per 10-week-old mouse (n = 20-26 per group). (**I**) Serum concentration of 4 kDa FITC–dextran over time after oral gavage in 10-week-old mice (n = 5 per group). *C. rodentium*-infected WT mice (9 days post-infection; n = 4) served as positive controls. (**J**) Quantitative RT–PCR analysis of transcripts related to $T_{FH}$ differentiation in total PP cells from indicated mice (n = 5-6 per group pooled from two experiments). (**K**) mRNA levels of antimicrobial peptides (AMPs) in intestinal tissues (n = 4 per group). Statistical analysis was performed using the unpaired Mann–Whitney U test or two-way ANOVA, as appropriate. Each dot represents one mouse.

**Figure S2**

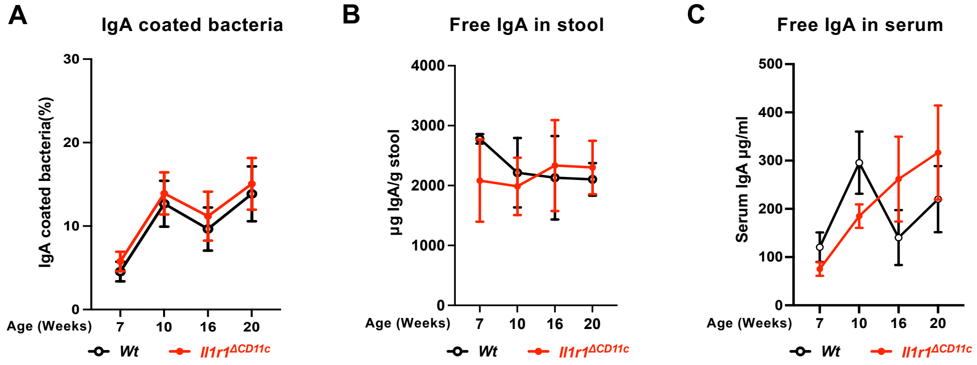


**Supplementary Figure 2. The heightened levels of antibody in the intestines of *Il1r1*-deficient mice at steady state were not depending on CD11c⁺ cells intrinsic IL-1R1 signaling.**
(**A)** the ratio of IgA-bound bacteria over time **(B and C**) Free IgA levels assessed by ELISA overtime in the stool (B) or the serum (C) of indicated mice (n=10 each group). Statistical analysis was performed using two-way ANOVA.

#### Figure S3

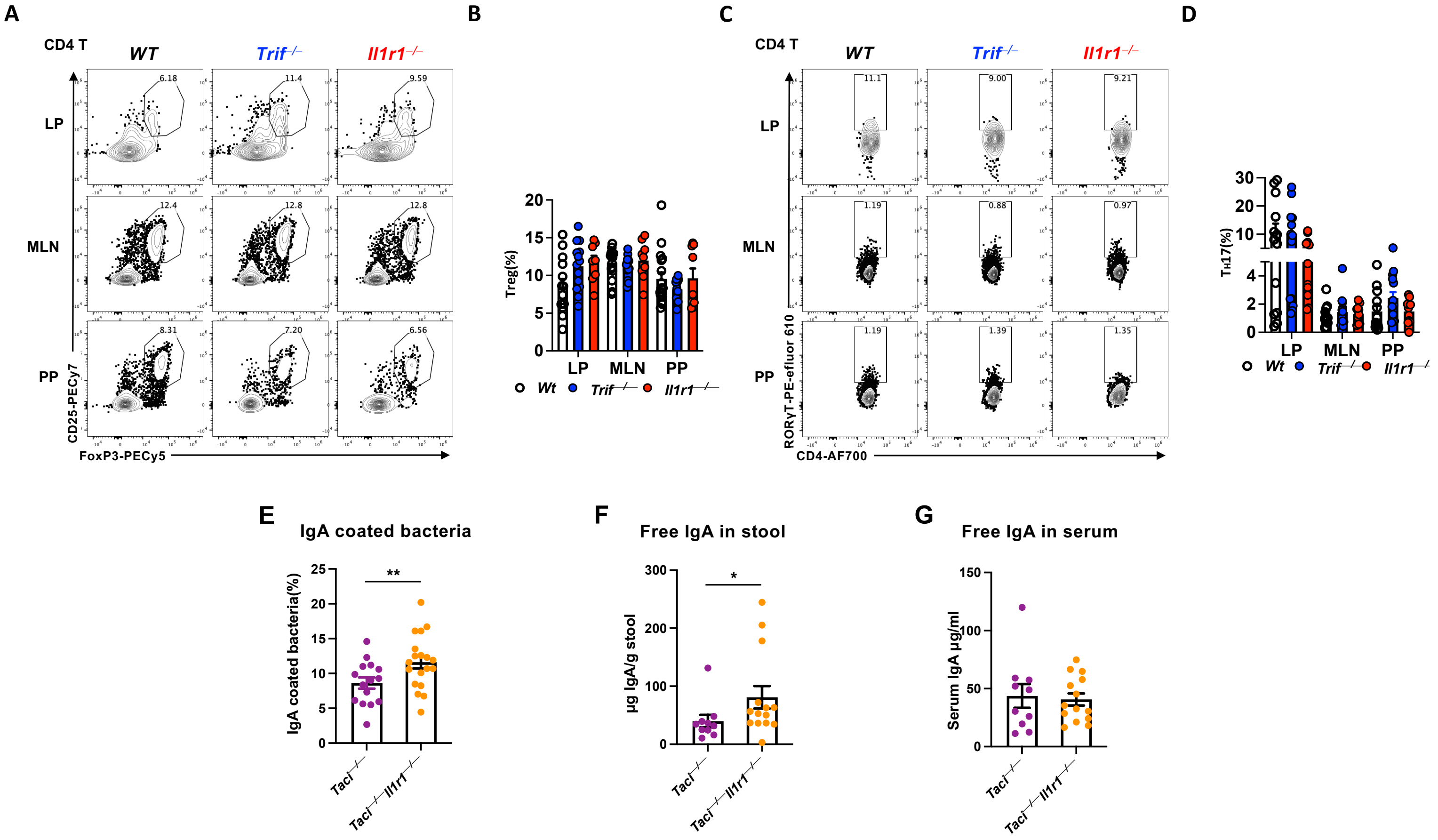


**<u>Supplementary Figure3</u>. *Il1r1* deficiency enhances intestinal IgA production independently of the T-independent pathway, while *Trif-* and *Il1r1-* deficiency do not alter Treg or $T_H17$ differentiation at steady state.**

(**A and B**) Dot plots (A) and percentages (B) of Foxp3$^+$ Treg cells among CD4 T cells. (**C and D**) Dot plots (C) and percentages (D) of RORγt$^+$ $T_H17$ cells among CD4 T cells. (n = 10-12 pooled from three experiments) (**E**) The ratio of IgA-bound bacteria of the 10 weeks old mice (**F and G**) IgA level in the stool (F) and the IgA level in the serum (G) of the 10 weeks old mice. n=15-19 pooled from three experiments. LP: Lamina Propria of the small intestines. MLN: Mesenteric Lymph nodes. PP: Peyer's patches. Statistical analysis was performed using unpaired Mann–Whitney U test. $p < 0.05$; $p < 0.01$; *$p < 0.001$. Each dot represents one mouse

Figure S4

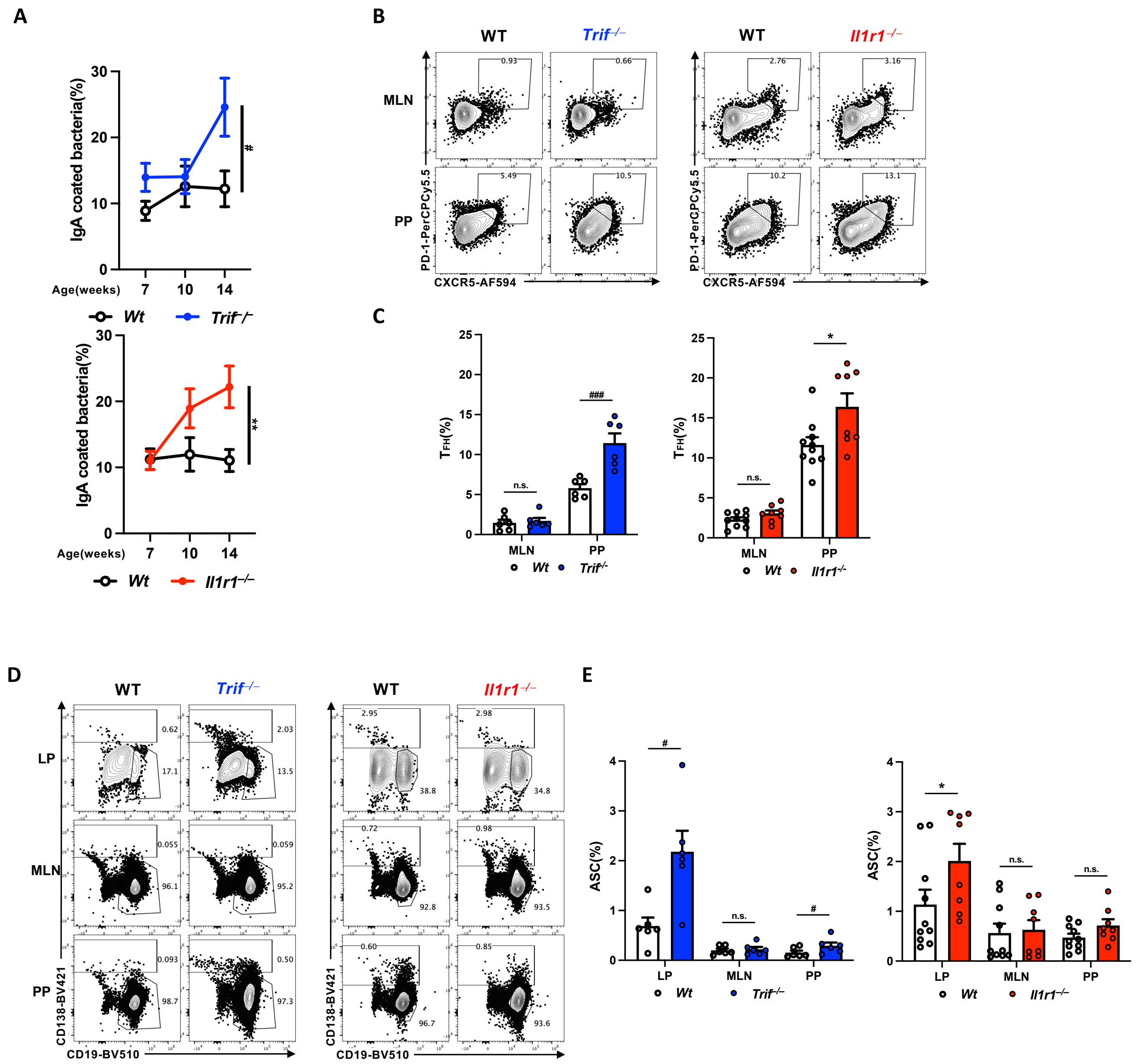


**Supplementary Figure 4. Enhanced IgA, B and T cell activation in the intestines of *Trif*$^{-/-}$ and *Il1r1*$^{-/-}$ mice compared to their respective littermate controls.**

**(A)** The ratio of IgA-bound bacteria over time implicated heightened levels of IgA in the intestines of *Trif*- and *Il1r1*-deficient littermate mice at steady state (n = 11-13 per group pooled from three independent experiments). (**B, C**) dot plots (B) and percentages (C) of $T_{FH}$ cells. (**D, E**) dot plots (D) and percentages (E) of CD138$^{+}$ cells (n = 6-10 per group pooled from two independent experiments). LP: Lamina Propria of the small intestines. MLN: Mesenteric Lymph nodes. PP: Peyer's patches. ASC: antibody secretion cells. Statistical analysis was performed using the unpaired Mann–Whitney U test or two-way ANOVA, as appropriate * p<0.05; ** p<0.01; *** p<0.001. #: WT to *Trif*$^{-/-}$, *: WT to *Il1r1*$^{-/-}$. Each dot represents one mouse.

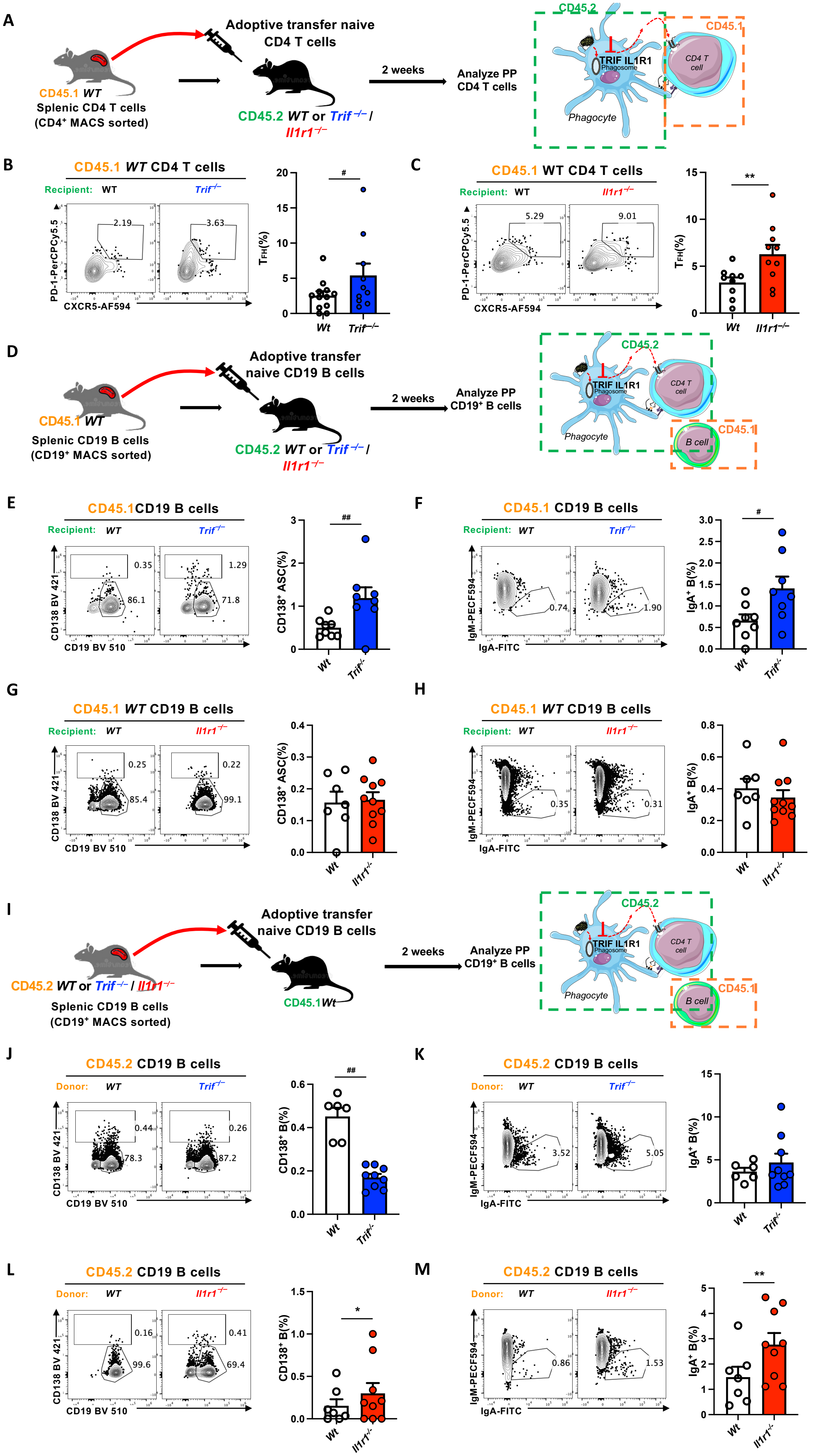
A
Adoptive transfer naive CD4 T cells
CD45.1 WT
Splenic CD4 T cells
(CD4⁺ MACS sorted)
CD45.2 WT or Trif −/− / Il1r1 −/−
2 weeks
Analyze PP CD4 T cells
CD45.2
CD45.1
TRIF IL1R1
Phagosome
CD4 T cell
Phagocyte
B
CD45.1 WT CD4 T cells
Recipient: WT Trif −/−
2.19
3.63
PD-1-PerCPCy5.5
CXCR5-AF594
TFH(%)
#
Wt Trif −/−
C
CD45.1 WT CD4 T cells
Recipient: WT Il1r1 −/−
5.29
9.01
PD-1-PerCPCy5.5
CXCR5-AF594
TFH(%)
**
Wt Il1r1 −/−
D
Adoptive transfer naive CD19 B cells
CD45.1 WT
Splenic CD19 B cells
(CD19⁺ MACS sorted)
CD45.2 WT or Trif −/− / Il1r1 −/−
2 weeks
Analyze PP CD19⁺ B cells
CD45.2
CD45.1
TRIF IL1R1
Phagosome
CD4 T cell
B cell
Phagocyte
E
CD45.1CD19 B cells
Recipient: WT Trif −/−
0.35
86.1
1.29
71.8
CD138 BV 421
CD19 BV 510
CD138⁺ ASC(%)
##
Wt Trif -/-
F
CD45.1 CD19 B cells
Recipient: WT Trif −/−
0.74
1.90
IgM-PECF594
IgA-FITC
IgA⁺ B(%)
#
Wt Trif -/-
G
CD45.1 WT CD19 B cells
Recipient: WT Il1r1 −/−
0.25
85.4
0.22
99.1
CD138 BV 421
CD19 BV 510
CD138⁺ ASC(%)
Wt Il1r1 -/-
H
CD45.1 WT CD19 B cells
Recipient: WT Il1r1 −/−
0.35
0.31
IgM-PECF594
IgA-FITC
IgA⁺ B(%)
Wt Il1r1 -/-
I
Adoptive transfer naive CD19 B cells
CD45.2 WT or Trif −/− / Il1r1 −/−
Splenic CD19 B cells
(CD19⁺ MACS sorted)
CD45.1Wt
2 weeks
Analyze PP CD19⁺ B cells
CD45.2
CD45.1
TRIF IL1R1
Phagosome
CD4 T cell
B cell
Phagocyte
J
CD45.2 CD19 B cells
Donor: WT Trif −/−
0.44
78.3
0.26
87.2
CD138 BV 421
CD19 BV 510
CD138⁺ B(%)
##
Wt Trif -/-
K
CD45.2 CD19 B cells
Donor: WT Trif −/−
3.52
5.05
IgM-PECF594
IgA-FITC
IgA⁺ B(%)
Wt Trif -/-
L
CD45.2 CD19 B cells
Donor: WT Il1r1 −/−
0.16
99.6
0.41
69.4
CD138 BV 421
CD19 BV 510
CD138⁺ B(%)
*
Wt Il1r1 -/-
M
CD45.2 CD19 B cells
Donor: WT Il1r1 −/−
0.86
1.53
IgM-PECF594
IgA-FITC
IgA⁺ B(%)
**
Wt Il1r1 -/-

**Figure 3. TRIF and IL-1R1 regulate $T_{FH}$ and B cell differentiation in Peyer's patches.**

(**A–C**) Adoptive transfer of $CD45.1^+$ WT CD4 T cells into $CD45.2^+$ WT, *Trif*$^{-/-}$(B), or *Il1r1*$^{-/-}$(C) recipients. (A) Experimental schematic. (B and C) Representative dot plots (left) and frequencies (right) of donor-derived PD-$1^+$CXCR$5^+$ $T_{FH}$ cells in Peyer's patches (PP) two weeks post-transfer. (**D–H**) WT $CD45.1^+CD19^+$ B cells were transferred into $CD45.2^+$ WT, *Trif*$^{-/-}$(E, F), or *Il1r1*$^{-/-}$ (G, H) recipients. (D) Schematic. (E and G) Percentages of donor-derived $CD138^+$ ASCs in PP. (F and H) Percentages of donor-derived $IgA^+$ B cells. (**I–M**) Reciprocal adoptive transfer of $CD45.2^+$ WT, *Trif*$^{-/-}$(J, K), or *I Il1r1*$^{-/-}$(L, M) $CD19^+$ B cells into $CD45.1^+$ WT recipients. (I) Schematic. (J and L) Frequencies of donor-derived $CD138^+$ ASCs. (K and M) Frequencies of donor-derived $IgA^+$ B cells. Statistical analysis: unpaired Mann–Whitney *U* test;* $p<0.05$; ** $p<0.01$; *** $p<0.001$. *: WT to *Il1r1*$^{-/-}$, #: WT to *Trif*$^{-/-}$. Each dot represents one mouse. (n = 4-9 per group pooled from two independent experiments)

#### Figure S5

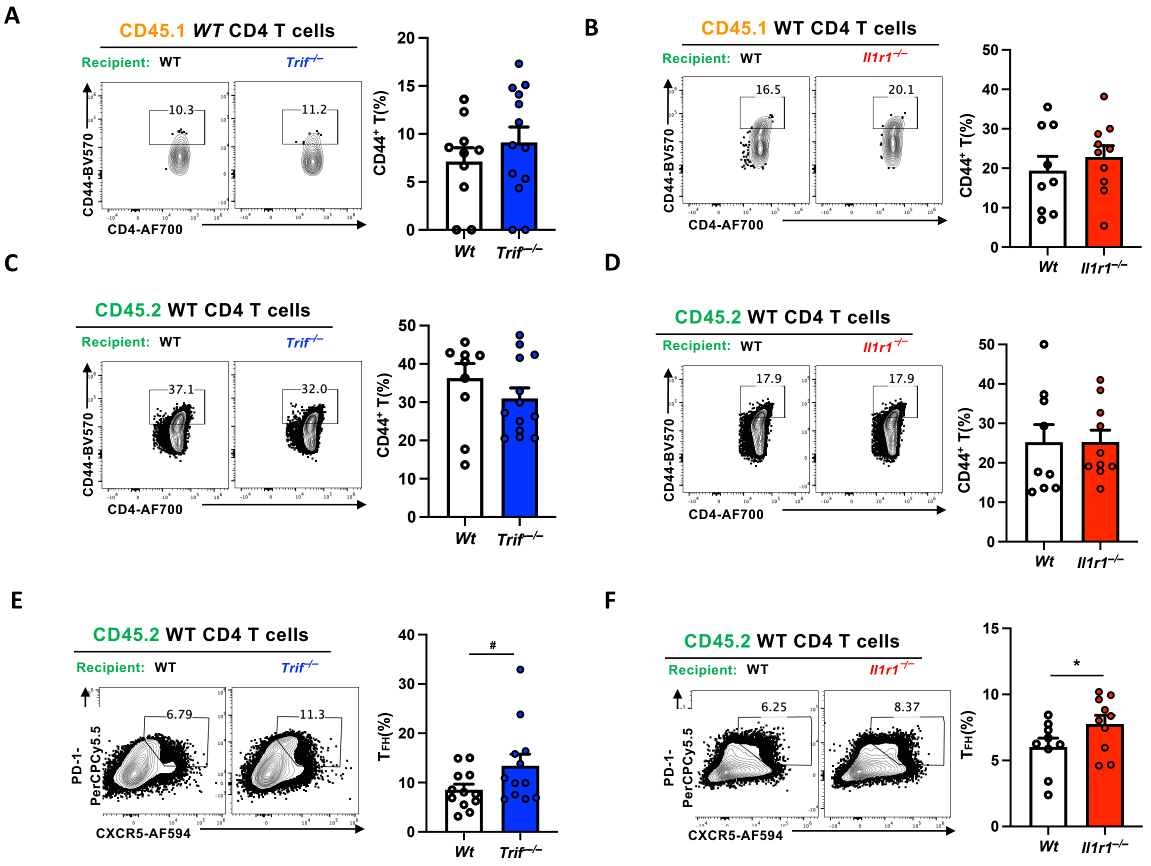


**Supplementary Figure 5. The *Trif*$^{-/-}$ and *Il1r1*$^{-/-}$ recipients mice showed higher ratio of PP $T_{FH}$ cells.** Experiments associated with Fig 3 A. (**A**) Representative dot plots (left) and percentages (right) of donor-derived $CD44^+$ CD4 T cells from Peyer's patches (PP) two weeks after adoptive transfer of $CD45.1^+$ WT CD4 T cells into $CD45.2^+$ WT or *Trif*$^{-/-}$,recipient mice. (**B**) Dot plots (left) and percentages (right) of donor-derived $CD44^+$ CD4 T cells from PP two weeks after adoptive transfer of $CD45.1^+$ WT CD4 T cells into $CD45.2^+$ WT or *Il1r1*$^{-/-}$ recipient mice. (**C**) Representative dot plots (left) and percentages (right) of recipient $CD44^+$ CD4 T cells from Peyer's patches (PP) two weeks after adoptive transfer of $CD45.1^+$ WT CD4 T cells into $CD45.2^+$ WT or *Trif*$^{-/-}$ recipient mice. (**D**) Dot plots (left) and percentages (right) of recipient $CD44^+$ CD4 T cells from PP two weeks after adoptive transfer of $CD45.1^+$ WT CD4 T cells into $CD45.2^+$ WT or *Il1r1*$^{-/-}$ recipient mice. (**E**) Dot plots (left) and percentages (right) of recipient PD-$1^+$$CXCR5^+$ $T_{FH}$ cells from PP in the same experimental setting as (C). (**F**) Dot plots (left) and percentages (right) of recipient PD-$1^+$$CXCR5^+$ $T_{FH}$ cells from PP in the same experimental setting as (D). Statistical analysis: Statistical analysis was performed using unpaired Mann–Whitney *U* test. $p < 0.05$; $p < 0.01$; *$p < 0.001$. #: WT to *Trif*$^{-/-}$, *: WT to *Il1r1*$^{-/-}$. Each dot represents one mouse (n = 8-12 per group pooled from two independent experiments).

**Figure S6**

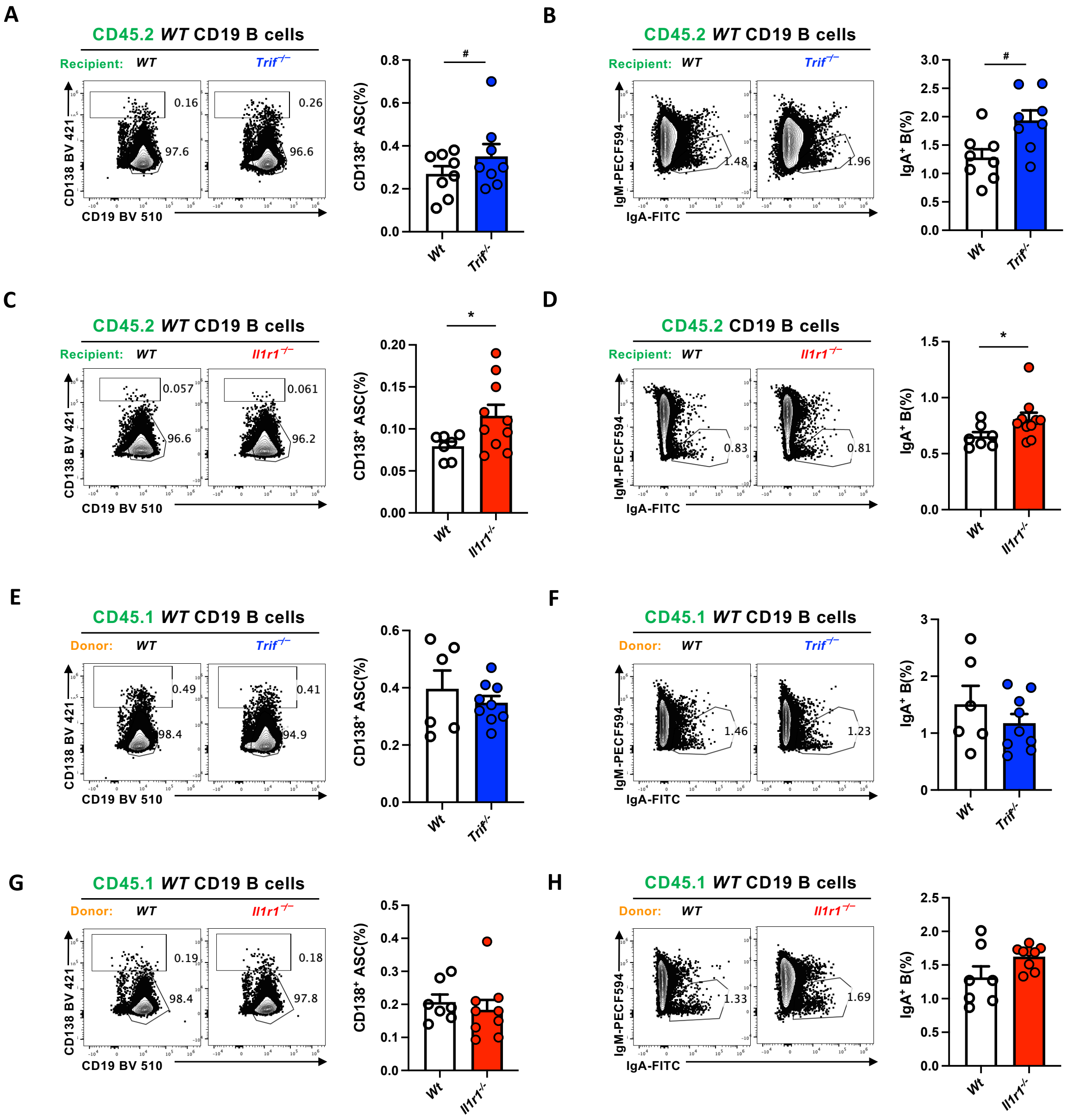


**Supplementary Figure 6. *Trif*$^{-/-}$ and *Il1r1*$^{-/-}$ recipients mice showed higher ratio of PP IgA$^{+}$ B cells.** (**A-D**) Experiments associated with Fig 3 D. (A) Representative dot plots (left) and percentages (right) of recipient CD138$^{+}$ ASC from Peyer's patches (PP) two weeks after adoptive transfer of CD45.2$^{+}$ WT or *Trif*$^{-/-}$ CD19$^{+}$ B cells into CD45.1$^{+}$ WT recipient mice. (B) Dot plots (left) and percentages (right) of recipient IgA$^{+}$ B cells from PP in the same experimental setting as (A). (C) Dot plots (left) and percentages (right) of recipient CD138$^{+}$ ASC cells from PP two weeks after adoptive transfer of CD45.1$^{+}$ WT CD19$^{+}$ B cells into CD45.2$^{+}$ WT or *Il1r1*$^{-/-}$ recipient mice. (D) Dot plots (left) and percentages (right) of recipient IgA$^{+}$ B cells cells from PP in the same experimental setting as (C). (**E-H**) Experiments associated with Fig 3 **I**. (E**)** Dot plots (left) and percentages (right) of recipient CD138$^{+}$ ASC cells from PP two weeks after adoptive transfer of CD45.2$^{+}$ WT or *Trif*$^{-/-}$ CD19$^{+}$ B cells into CD45.1$^{+}$ WT recipient mice. (F) Dot plots (left) and percentages (right) of recipient IgA$^{+}$ B cells cells from PP in the same experimental setting as (E). (G) Dot plots (left) and percentages (right) of recipient CD138$^{+}$ ASC cells from PP two weeks after adoptive transfer of CD45.2$^{+}$ WT or *Il1r1*$^{-/-}$ CD19$^{+}$ B cells into CD45.1$^{+}$ WT recipient mice. (H) Dot plots (left) and percentages (right) of recipient IgA$^{+}$ B cells cells from PP in the same experimental setting as (G). Statistical analysis: unpaired Mann–Whitney *U* test;* p<0.05; ** p<0.01; *** p<0.001. *: WT to *Il1r1*$^{-/-}$, #: WT to *Trif*$^{-/-}$. Each dot represents one mouse (n = 7-13 per group pooled from two independent experiments).

#### Figure S7

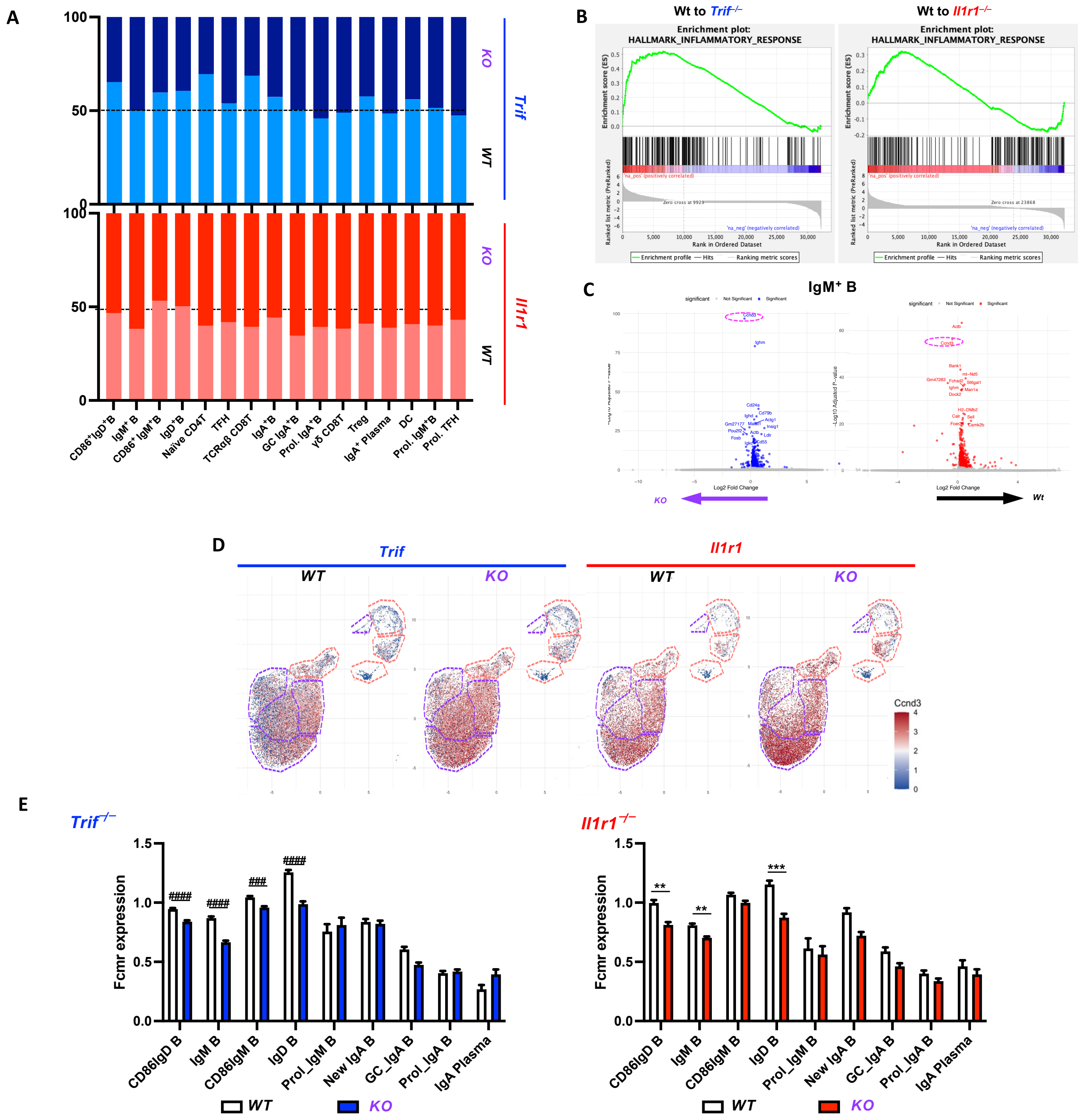


**Supplementary Figure 7. Increased immune activation programs in Peyer's patches of *Trif*$^{-/-}$ and *Il1r1*$^{-/-}$ mice compared to their littermate controls.**

(**A**) A stacked bar chart was used to depict the relative proportions of identified cell types across different mouse strains (Top: *Trif*$^{-/-}$ ; bottom: *Il1r1*$^{-/-}$ mice comparing to WT littermate control). (**B**) Representative GSEA plot showing enrichment of the "inflammatory response" gene set in DCs from *Trif*$^{-/-}$ and *Il1r1*$^{-/-}$ mice compared to *WT*. (**C**) Volcano plot of differentially expressed genes in IgM+ B cells; *Ccnd3* is highlighted with a pink dashed circle. (**D**) UMAP plot of *Ccnd3* expression across B cell clusters. (**E**) Bar plots of *Fcmr* expression in indicated clusters for *Trif*$^{-/-}$ (left) and *Il1r1*$^{-/-}$(right) mice compared to *WT*. (Statistical analysis: unpaired Mann–Whitney *U* test; * p<0.05; ** p<0.01; *** p<0.001. #: WT to *Trif*$^{-/-}$, *: WT to *Il1r1*$^{-/-}$.

#### Figure S8

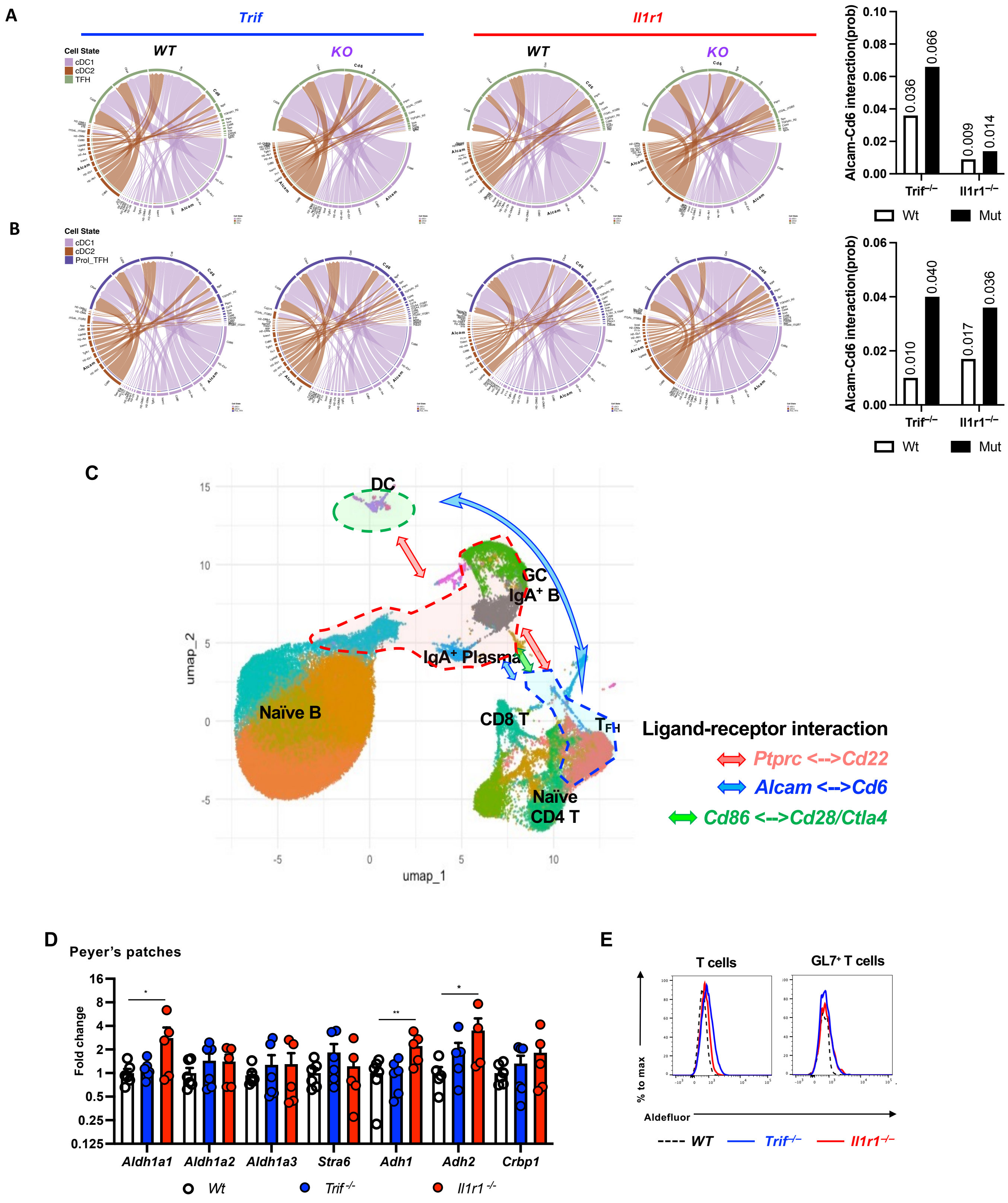


**Supplementary Figure 8. Shift in cell-cell interaction within Peyer's patches of *Trif*$^{-/-}$ and *Il1r1*$^{-/-}$ mice compared to their littermate controls.**

(**A, B**) CellChat-derived chord diagrams of ligand–receptor interactions among cDC1, cDC2, $T_{FH}$ cells (A), and proliferating $T_{FH}$ cells (B) in the PP. (**C**) Schematic illustration of interactions between different cell types. (**D**) Quantitative RT–PCR analysis of transcripts related to retinoic acid metabolism in total PP cells from indicated mice (n = 5-6 per group pooled from two experiments). (**E**) Aldefluor histograms of *Trif*$^{-/-}$ and *Il1r1*$^{-/-}$ mice compared to WT, separated by T cell subtype. Statistical analysis: unpaired Mann–Whitney *U* test; * p<0.05; ** p<0.01; *** p<0.001. #: WT to *Trif*$^{-/-}$, *: WT to *Il1r1*$^{-/-}$.

**Figure S9**

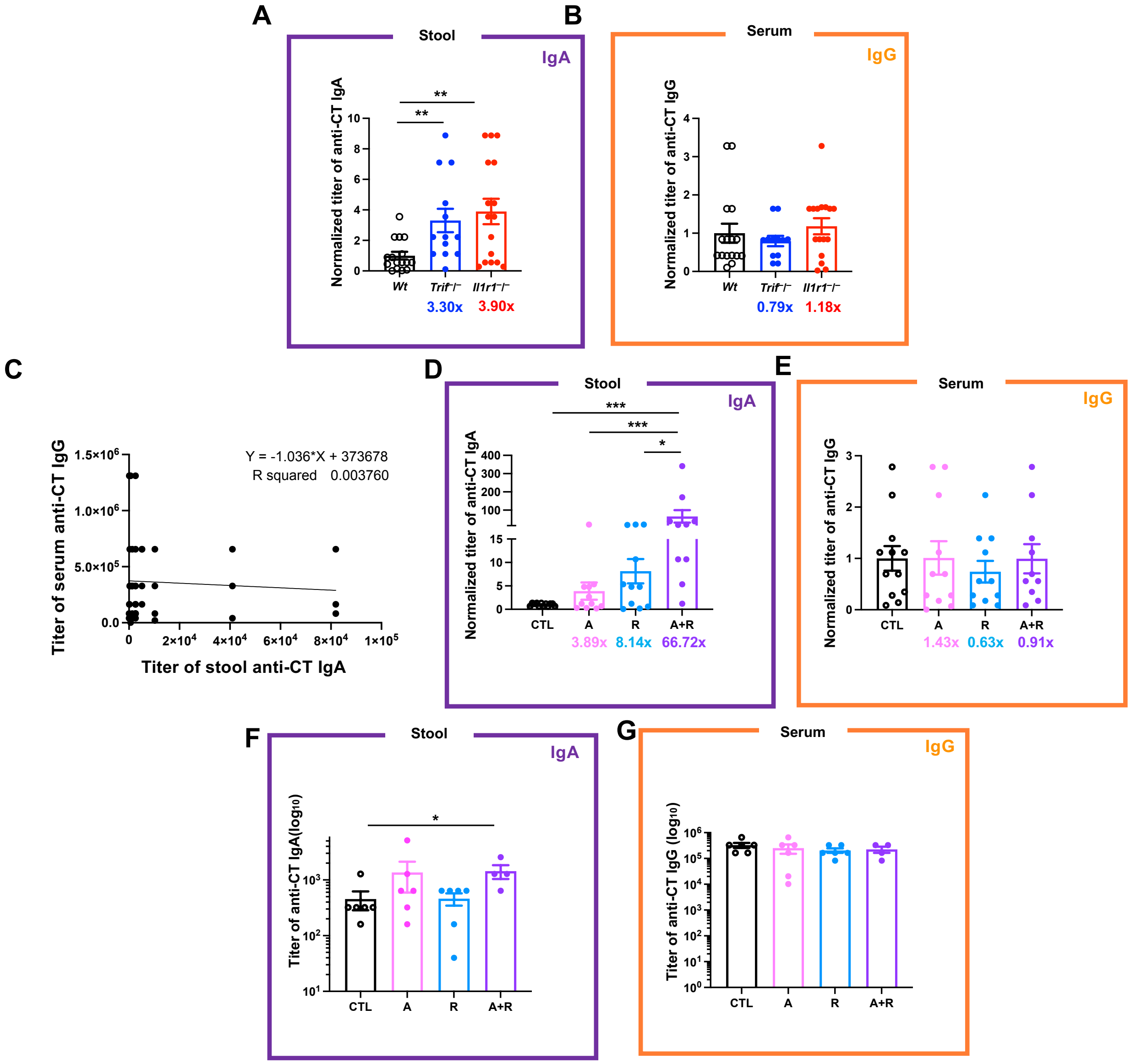


**<u>Supplementary Figure 9</u>. Related to Figure 5. Additional analyses of mucosal antigen-specific IgA responses following cholera toxin vaccination.**

(**A–B**) Related to Figure 5A. (A) Levels of stool CT-specific IgA, normalized to the mean of the control group (*WT*) in each experiment, from the indicated mice 4 weeks after the first vaccination. (B) Normalized serum CT-specific IgG levels in the indicated mice 4 weeks after the first vaccination. (**C–E**) Related to Figure 5D. (C) Scatter plot of stool anti-CT IgA titers versus serum anti-CT IgG titers. Each dot represents one mouse. (D**)** Normalized stool CT-specific IgA levels in the indicated mice 6 weeks after the first vaccination. (E) Normalized serum CT-specific IgG levels in the indicated mice 6 weeks after the first vaccination. (**F)** Normalized stool CT-specific IgA levels in the indicated mice, 1 weeks after the re-immunization. (**G)** Normalized serum CT-specific IgG levels in the indicated mice, 1 weeks after the re-immunization. CT: cholera toxin; CTL: control; A: Anakinra; R: ATRA; A+R: Anakinra plus ATRA; WPV: weeks after the first vaccination. Statistical analysis was performed using unpaired Mann–Whitney *U* test. $p < 0.05$; $p < 0.01$; *$p < 0.001$. Each dot represents one mouse.

#### Figure 10

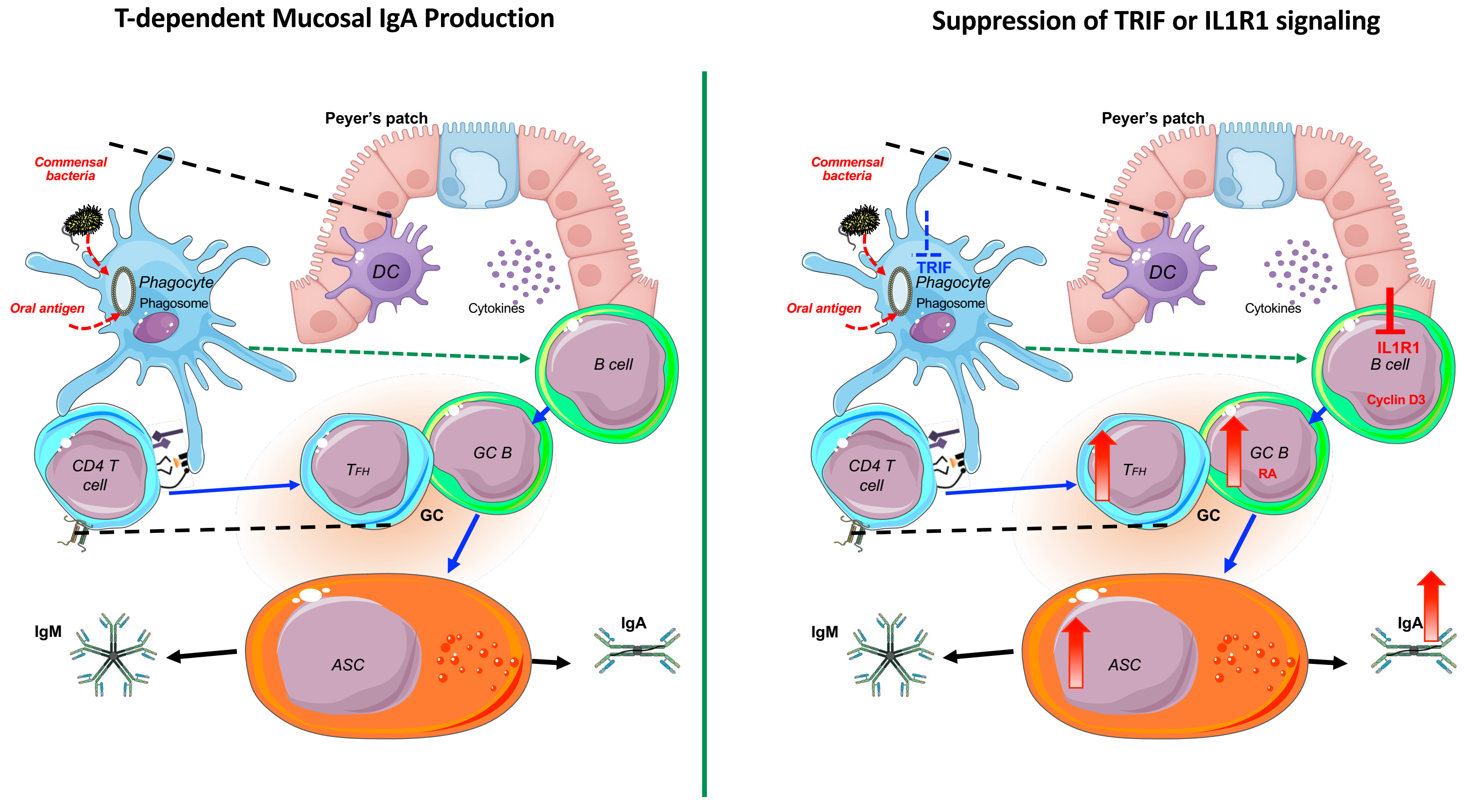


**<u>Supplementary Figure 10</u>. Proposed Model for Enhanced Mucosal IgA Responses Following Suppression of TRIF- and IL-1R1-Associated Signaling.**
Genetic loss of *Trif* or *Il1r1*, or pharmacologic IL-1R1 blockade with Anakinra, releases inhibitory constraints on Peyer's patch IgA responses. In this model, IL-1R1 acts, at least in part, as a B cell–intrinsic brake, whereas TRIF- and IL-1R1-associated pathways collectively influence the Peyer's patch microenvironment. This process is associated with altered RA-related activity and increased *Ccnd3*/Cyclin D3 expression, promoting GC-associated B cell differentiation, ASC generation, and intestinal IgA production. Adapted from Servier Medical Art, licensed under CC BY 4.0.
